# Hypothalamic asymmetry in hemisphere-specific neuroendocrine signaling

**DOI:** 10.64898/2026.02.16.705997

**Authors:** Hiroyuki Watanabe, Yaromir Kobikov, Olga Nosova, Alisa Lukoyanova, Daniil Sarkisyan, Emma Lindström, Vladimir Galatenko, Yoichi Ueta, Takashi Maruyama, Alfhild Grönbladh, Mathias Hallberg, Jens Schouenborg, Rui Francisco Ferreira, Susana Maria Silva, Nikolay Lukoyanov, Igor Lavrov, Mengliang Zhang, Georgy Bakalkin

**Affiliations:** Department of Pharmaceutical Biosciences, Uppsala University, Uppsala, Sweden; Research Affiliate, Department of Pharmaceutical Biosciences, Uppsala University. Present address: Tashkent, Uzbekistan; Department of Immunology, Genetics and Pathology and Science for Life Laboratory, Uppsala University, Uppsala, Sweden; Evotec International GmbH, Göttingen, Germany; Neuronal Networks Group, i3S-Institute for Research and Innovation in Health, University of Porto, Porto, Portugal; Department of Physiology, School of Medicine, University of Occupational and Environmental Health, Japan; Neuronano Research Center, Department of Experimental Medical Science, Lund University, Lund, Sweden; Master’s course in Biochemistry, Faculty of Sciences, University of Porto, Porto, Portugal; Unit of Anatomy, Department of Biomedicine, Faculty of Medicine, University of Porto, Porto, Portugal; CINTESIS@RISE, Porto, Portugal; Department of Neurology, Mayo Clinic, Rochester, MN, USA; Department of Molecular Medicine, University of Southern Denmark, Odense, Denmark

**Keywords:** neuropeptides, Arg-vasopressin, hypothalamic asymmetry, endocrine system, neural circuit, DREADD, gene co-expression networks

## Abstract

Brain lesions classically cause contralateral sensorimotor and postural deficits, attributed to the decussation of descending neural pathways. However, recent findings reveal that, beyond neural mechanisms, contralateral effects can also be mediated by the neuroendocrine system via humoral pathways. This raises the possibility that the brain regulates left- and right-sided peripheral processes through hypothalamic neurohormones released into the bloodstream. For such spatially targeted endocrine signaling to occur, hemisphere-specific neural activity must be encoded into side-specific hormonal output—requiring a lateralized organization of hypothalamic neuroendocrine systems. Here, we report molecular asymmetries in the rat hypothalamus that support this mechanism. Transcriptomic analysis revealed asymmetric expression of eleven neurohormonal genes, including *Gnrh1*, *Cck*, and *Trh*, along with distinct left-right side-specific gene co-expression networks. Chemogenetic stimulation of Arg-vasopressin neurons in vasopressin-hM3Dq-mCherry transgenic rats produced generalized changes in these networks—predominantly in the right hypothalamus—suggesting that hypothalamic neurohormonal circuits function as integrated, lateralized ensembles. Stereological analysis revealed asymmetric coordination of vasopressin neurons within the paraventricular nucleus, with the left rostral region decoupled from the right rostral and caudal subregions. Functionally, gonadotropin-releasing hormone, cholecystokinin-8, and thyrotropin-releasing hormone—administered intracisternally in rats with complete spinal cord transection—elicited side-specific peripheral responses, measured as hindlimb postural asymmetry in a binary left–right output model. These neurohormonal effects were therefore transmitted via the humoral route. These findings suggest that multiple hypothalamic neurohormones and their integrated networks are asymmetrically organized, and that their lateralization may be necessary for hemisphere-specific hormonal regulation of peripheral systems.

## INTRODUCTION

Traumatic brain injury and stroke in patients, and brain lesions in animal experiments cause contralateral sensorimotor and postural deficits including asymmetric posture and reflexes (1–5). According to the neurology dogma contralateral deficits result from the decussation of the descending neural pathways (6–13). Beyond neural mechanism, the contralateral injury effects may be mediated by the neuroendocrine system through the humoral pathway (6,14–17). Side-specific endocrine signaling has been identified in animals in which the spinal cords were completely transected before the brain was unilaterally lesioned. In these animals, injury to the hindlimb sensorimotor cortex elicited hindlimb postural asymmetry (HL-PA), a binary response involving flexion of the contralateral hindlimb, as well as asymmetries in withdrawal reflexes. Serum from rats with unilateral brain injury (UBI) induced HL-PA in naïve animals, while hypophysectomy abolished UBI-induced asymmetries in spinalized rats. Two neurohormones, Arg-vasopressin (AVP) and β-endorphin, were identified as mediators of left-brain injury effects eliciting right hindlimb responses (14). These neurohormones also induced right-side motor responses in animals with intact brain. Complementary, dynorphin and Met-enkephalin appear to mediate right-brain injury effects via the humoral pathway, resulting in left hindlimb flexion (17). Thus, the molecular endocrine signals elicited by the hypothalamic-pituitary system, convey information on the side of brain injury.

This topographic, left–right-specific neuroendocrine system (T-NES) may function through i) *encoding* of hemisphere-specific neural signals into side-specific neurohormones by the hypothalamus and pituitary, ii) transmission of these signals via the bloodstream, and iii) *decoding* them at peripheral targets, such as the spinal cord, into lateralized physiological and endocrine responses (6,14,17). The differential encoding of neural signals from the two anatomically symmetric hemispheres into side-specific neurohormonal messages may rely on a lateralized, hemisphere-specific organization of the T-NES, primarily the hypothalamus—the interface between the nervous and endocrine systems (14,17). Ipsilateral inputs from the hemispheres may selectively activate neurohormones with side-specific actions, if those neurohormones are asymmetrically organized in the hypothalamus. Neurohormones lateralized to the left may preferentially respond to left-hemisphere signals, while those lateralized to the right may be more responsive to right-hemisphere input. Once released into circulation, these neurohormones may exert differential effects on the left and right sides of the body acting directly or by stimulating secretion of pituitary hormones with side-specific actions (14,17).

Our previous study identified asymmetric expression of three neurohormonal genes— *Pomc*, *Trh*, and *Ghrh*—in the rat hypothalamus (17). In addition, both neurohormonal and neuroplasticity-related genes showed side-specific patterns of co-expression, which could be classified into two distinct gene co-expression networks: a left-dominant network (LdN) and a right-dominant network (RdN). However, these findings were derived from a neuropathological model involving deep anesthesia, complete spinal cord transection, and unilateral brain injury, and were based on a relatively limited set of neurohormonal genes. Therefore, the generality of this lateralized gene expression architecture remains to be validated under normal physiological conditions and across other experimental models.

In the present study, we first validated the lateralized expression of individual neurohormonal genes and their co-expression networks using three independent rat cohorts (described below), including naïve animals. To expand beyond our prior findings, we analyzed a broader set of genes encompassing the majority of hypothalamic neurohormones. Second, recognizing that neurohormonal genes are expressed across distinct neuronal populations and subregions of the hypothalamus (18,19), we reasoned that their co-expression patterns likely reflect functional interactions between neuroendocrine circuits. Under this hypothesis, perturbation of a single neurohormonal pathway would alter the activity of other, interconnected circuits, manifesting as coordinated changes in gene expression and co-expression. To test this, we analyzed how chemogenetic stimulation of AVP-expressing hypothalamic neurons—a lateralized neurohormone known to induce side-specific effects (14)—influences the expression and co-expression of other neurohormonal genes, with a specific focus on left–right asymmetries. Third, we hypothesized that lateralized hypothalamic neurohormones may initiate side-specific signaling within the T-NES by stimulating the release of humoral messengers into the circulation that convey hemisphere-specific information to peripheral targets or may themselves function as such messengers. To investigate this, we centrally administered three lateralized hypothalamic neurohormones and assessed their ability to elicit peripheral effects in rats with disrupted descending neural pathways. Finally, to explore whether AVP itself is anatomically lateralized, we examined its distribution within the paraventricular nucleus (PVN) using stereology-based quantitative immunohistochemistry.

To address these aims, we employed three rat cohorts: (i) naïve Wistar rats; (ii) transgenic rats with chemogenetically activated AVP neurons; and (iii) rats with complete thoracic spinal cord transection, used to assess hypothalamic–spinal coordination in the absence of descending neural control (for study design, see **Supplementary Figure 1**). In the latter cohort, the effects of unilateral ablation of the sensorimotor cortex on endocrine signaling were evaluated.

Our findings demonstrate that a subset of hypothalamic neurohormonal genes is asymmetrically expressed, and suggest that neuroendocrine neurons are functionally interconnected and organized into lateralized co-expression networks. Notably, each of the three lateralized neurohormones tested was sufficient to induce side-specific peripheral effects via the general circulation. Together, these results support the concept that lateralized neuropeptidergic systems are integral components of a topographic encoding mechanism, by which the hypothalamus transforms hemisphere-specific neural signals into spatially resolved endocrine outputs.

Because definitions of peptide neurohormones and neuropeptides often overlap, we use the term *neurohormones* throughout to include all peptide regulators acting in this system. Abbreviations and terms are listed in the glossary.

## MATERIALS AND METHODS

The methods used in this study, including the UBI model, surgeries, gene expression analysis, neuroplasticity-related genes analyzed, analysis of HL-PA and resistance to stretch, and statistical analysis of physiological and molecular data, have been described in our previous papers (14,17).

### Animals, surgery and treatments

Male Wistar rats (Janvier-labs, France) weighing 190-250 g were used as a cohort of naïve rats in molecular studies (n = 11) and for physiological analysis of neurohormonal effects (n = 41). For molecular studies, rats were euthanized between 10:00 and 12:00 AM. The second cohort consisted of transgenic Wistar rats expressing a human muscarinic acetylcholine receptor (hM3Dq) under the AVP promoter (AVP-hM3Dq-mCherry rats: 12 males and 10 females weighing 190-250 g). The AVP-hM3Dq-mCherry heterozygous transgenic rats were produced at University of Occupational and Environmental Health, Kitakyushu, Japan (20) and rederivation was performed by Janvier Labs (France). Further breeding of the heterozygous transgenic rats was performed by pairing the wild type and transgenic Wistar rats at Lund University (Sweden). Genotyping was performed by PCR analysis of genomic DNA extracted from rat ear using primers: sense sequence, 5′-CAC CAT CTT CTT CAA GGA CGA C-3′; antisense sequence, 5′-ATG ATA TAG ACG TTG TGG CTG TTG T-3′) (21). The AVP-hM3Dq-mCherry rats were treated with either Clozapine-N-oxide dihydrochloride (CNO; 1 mg/mL/kg; 6 males and 5 females) to chemogenetically stimulate the Avp-neurons in the hypothalamus, or vehicle (2% DMSO in saline, 1 mL/kg; 6 males and 5 females) and sacrificed 3 h later. The animals of both cohorts received food and water ad libitum and were maintained in a 12-h day-night cycle (light on from 10:00 p.m. to 10:00 a.m.) at a constant environmental temperature of 21°C (humidity: 65%). Ethical approval was obtained from the Malmö/Lund ethical committee of animal experiments (Dnr. 5.8.18-17317/2021).

The third cohort consisted of male Wistar Hannover rats (Charles River Laboratories, Spain) with body weight of 150-200 g used for molecular analysis. Complete transection of the spinal cord was performed at the T2-T4 level and followed by left UBI (n = 12 rats) or left sham surgery (n = 11 rats). Tissue samples were collected five hours after initial spinal transection surgery or three hours after UBI / sham surgery. The animals received food and water *ad libitum* and were kept at a 12-h day-night cycle (light on from 07:00 a.m. to 07:00 p.m.) at a constant environmental temperature of 21°C (humidity: 65%). Approval for animal experiments was obtained from the ethical committee of the Faculty of Medicine of Porto University and Portuguese Direção-Geral de Alimentação e Veterinária (No. 0421/000/000/2018).

Additional cohort of male Wistar Hannover rats (Charles River Laboratories) was used for immunohistochemical experiments. It consisted of naive (n = 6), left UBI (n = 5), right UBI (n = 5) and sham-surgery (n = 5) groups. Naive rats were handled and treated in the same way as the other groups, but did not undergo surgery. Rats in the UBI groups underwent surgery to ablate either the left or right hindlimb sensorimotor cortex as described below. Rats in the sham surgery group received treatments identical to the UBI groups, including opening the skull, but cortical tissue was not aspirated. Brain tissue was collected 3-5 days after UBI or sham surgery.

### Spinal cord transection

The animals were anesthetized with sodium pentobarbital (intraperitoneal, I.P.; 40 mg/kg body weight, as an initial dose and then 6 mg/kg every hour; third rat cohort) or isoflurane (for treatment with neurohormones) anesthesia. Core temperature of the animals was controlled using a feedback-regulated heating system. Anaesthetized animals were mounted onto the stereotaxic frame and the skin of the back was incised along the midline at the level of the superior thoracic vertebrae. After laminectomy, a 3-4-mm spinal cord segment at the T2-T4 level was dissected and removed (14). The completeness of the transection was confirmed by (i) inspecting the cord during the operation to ensure that no spared fibers bridged the transection site and that the rostral and caudal stumps of the spinal cord were completely retracted; and (ii) examining the spinal cord in all animals after termination of the experiment.

### Brain surgery

UBI and sham surgery were preceded by a complete spinal cord transection. The head of the rats under pentobarbital anesthesia was mounted onto the stereotaxic frame and fixed in a position in which the bregma and lambda were located at the same horizontal level. After local injection of lidocaine (Xylocaine, 3.5 mg/ml) with adrenaline (2.2 μg/ml), the scalp was cut, open and a piece of the parietal bone located 0.5 – 4.0 mm posterior to the bregma and 1.8 – 3.8 mm lateral to the midline was removed (22). The part of the cerebral cortex located below the opening that includes the hind-limb representation area of the sensorimotor cortex was aspirated with a glass pipette (tip diameter 0.5 mm) connected to an electrical suction machine (Craft Duo-Vec Suction unit, Rocket Medical Plc, UK). Care was taken to avoid damaging the white matter below the cortex. After the ablation, bleeding was stopped with a piece of Spongostone and the bone opening was covered with a piece of TissuDura (Baxter, Germany). For sham operations, animals underwent the same anesthesia and surgical procedures, but the cortex was not ablated. After completion of all surgical procedures, the wounds were closed with 3-0 suture (AgnTho’s, Sweden) and the rat was kept under an infrared radiation lamp to maintain appropriate body temperature.

### Neurohormonal treatment design

The spinal cords of intact rats were completely transected at the T2-T4 level under isoflurane anesthesia and neurohormones or saline were administered into the cisterna magna (intracisternal route; 5 microliters/rat) (23,24). HL-PA and hindlimb resistance to stretch were analyzed before the injection, and 1 and 3 h after injection while the animals remained under anesthesia.

Gonadotropin-releasing hormone (GnRH) was dissolved in saline and injected at doses 0.1, 1, and 50 ng/rat. Cholecystokinin-8 (CCK-8) was dissolved in 80% DMSO at concentration 1 mg/ml, and a solution was diluted with saline immediately before injection at doses 0.1, 1, 10 and 30 ng/rat. Thyrotropin-releasing hormone (TRH) was dissolved in saline and injected at dose 1 ng/rat.

### Analysis of gene expression

Gene expression was analyzed in the left and right halves of the hypothalamus separately in three cohorts, and the lumbar spinal cord in the UBI-sham surgery cohort. For the analyses, hypothalamic tissue was taken from the area between the plane drawn through the anterior commissure and the optic chiasm and the vertical plane drawn through the posterior border of the mammillary body. The samples were snap frozen and stored at −80 °C until assay. The rats in the naïve, transgenic, and UBI-sham-surgery cohorts were euthanized and their brains were dissected between 1:00 p.m. and 4:00 p.m., 1:00 p.m. and 4:00 p.m. (3 h after treatment with CNO or vehicle), and 3:00 p.m. and 5:00 p.m. (5 h after spinalization), respectively.

### Quantitative real-time PCR

Total RNA was purified by using RNeasy Lipid Tissue Mini Kit (Qiagen, Valencia, CA, USA). RNA concentrations were measured with Nanodrop (Nanodrop Technologies, Wilmington, DE, USA). RNA (500 ng) was reverse-transcribed to cDNA with the cDNA iScript kit (Bio-Rad Laboratories, CA, USA) according to manufacturer’s protocol. cDNA samples were aliquoted and stored at -20°C. cDNAs were mixed with PrimePCR™ Probe assay and iTaq Universal Probes supermix (Bio-Rad) for qPCR with a CFX384 Touch™ Real-Time PCR Detection System (Bio-Rad Laboratories, CA, USA) according to manufacturer’s instructions. TagMan assay was performed in 384-well format with TagMan probes that are listed in **Supplementary Tables 1 and 2**.

All procedures were conducted strictly in accordance with the established guidelines for the qRCR based analysis of gene expression, consistent with the minimum information for publication of quantitative real-time PCR experiments guidelines (25,26). The raw qPCR data were obtained by the CFX Maestro™ Software for CFX384 Touch™ Real-Time PCR Detection System (Bio-Rad Laboratories, CA, USA). The mRNA levels of genes of interest were normalized to the geometric mean of expression levels of two reference genes *Actb* and *Gapdh*. GeNorm software was used to analyze the gene expression stabilities (M value) of the ten candidate reference genes (*Actb, B2m, Gapdh, Gusb, Hprt, Pgk, Ppia, Rplpo13a, Tbp*, and *Tfrc*). The calculation of the M value was based on the pairwise variation between two reference genes. If the M value was less than 1.5, it could be considered as a suitable reference gene. The smaller the M value, the higher the stability of gene expression levels (https://genorm.cmgg.be/ and (27)). The expression stability of candidate reference genes was computed for all sets of samples and identified *Actb* and *Gapdh* as the most stably expressed genes. For both regions analyzed, the gene expression stability (M values) did not exceed 0.5. The optimal number of reference genes was determined by calculating pairwise variation (V value) by GeNorm program. The V value for *Actb* and *Gapdh*, the top reference genes, was 0.12 that did not exceed the 0.15 threshold demonstrating that analysis of these two genes is sufficient for normalization.

### Hormonal, neurohormonal and neuroplasticity-related genes

In the hypothalamus the hypothalamic neurohormone, neuropeptide genes and their receptor genes including the arginine vasopressin *Avp*, cholecystokinin *Cck*, corticotropin releasing hormone *Crh*, galanin *Gal*, growth hormone releasing hormone *Ghrh*, ghrelin *Ghrl*, gonadotropin releasing hormone 1 *Gnrh1*, hypocretin neuropeptide precursor *Hert*, kiss-1 metastasis suppressor *Kiss*, neuropeptide Y *Npy*, neurotensin *Nts*, opioid receptor delta 1 *Oprd1*, opioid receptor kappa 1 *Oprk1*, opioid receptor mu 1 *Oprm1*, oxytocin *Oxt*, prodynorphin *Pdyn*, proenkephalin *Penk*, pro-melanin concentrating hormone *Pmch*, prepronociceptin *Pnoc*, proopiomelanocortin *Pomc*, prolactin releasing hormone *Prlh*, somatostatin *Sst*, thyrotropin releasing hormone *Trh*, urocortin 3 *Ucn3* and vasoactive intestinal peptide*, Vip* genes, and genes of the RAS system including the angiotensin I converting enzyme *Ace*, angiotensin converting enzyme 2 *Ace2*, angiotensinogen *Agt*, angiotensin II receptor type 1a*, Agtr1a*, angiotensin II receptor type 1b *Agtr1b*, angiotensin II receptor type 2 *Agtr2*, angiotensin II receptor associated protein *Agtrap*, Mas1 proto-oncogene G-protein-coupled receptor *Mas1 and* renin *Ren,* genes, were analyzed (**Supplementary Table 1**).

Genes selected as neuroplasticity-related were identified as such in several major studies. The selection of each gene from this set is justified by referring to these studies (see references below) and is not biased. To note there is no established view on how to categorize genes as neuroplasticity-related, and there are no lists of neuroplasticity-related genes consistent among studies. Thus, such a selection is arbitrary and a set of selected genes could not be comprehensive. The selected neuroplasticity-related genes were: *Arc*, activity-regulated cytoskeletal gene implicated in numerous plasticity paradigms; *Bdnf*, brain-derived neurotrophic factor regulating synaptogenesis; *cFos*, a neuronal activity dependent transcription factor; *Dlg4* gene encoding PSD95 involved in AMPA receptor-mediated synaptic plasticity and post NMDA receptor activation events; *Egr1* regulating transcription of growth factors, DNA damage, and ischemia genes; *Gap-43* coding for growth-associated protein Gap-43 that regulates axonal growth and neural network formation; *GluR1* and *Grin2b* coding for the glutamate ionotropic receptor AMPA Type Subunit 1 and NMDA receptor subunit, respectively, both involved in glutamate signaling and synaptic plasticity; *Grin2a* subunit of the glutamate receptors that regulates formation of neural circuits and their plasticity; *Homer-1* giving rise to homer scaffold protein 1, a component of glutamate signaling involved in nociceptive plasticity; *Pcsk6* gene encoding proprotein convertase subtilisin/kexin type 6 involved in post-translational modification; *Nfkbia* (I-Kappa-B-Alpha) that inhibits NF-kappa-B/REL complexes regulating activity-dependent inhibitory and excitatory neuronal function; *Syt4* (Synaptotagmin 4) playing a role in dendrite formation and synaptic growth and plasticity; and *Tgfb1* that gives rise to transforming growth factor β1 regulating inflammation, expression of neuropeptides and glutamate neurotoxicity (28–43) (**Supplementary Table 1**). The 47 hypothalamic genes were analyzed in the naïve Wistar and transgenic cohorts (*Agtr2* was excluded due to technical reasons) and 48 genes in the UBI-sham surgery cohort.

Expression data for the neuroplasticity-related genes along with the neuropeptide and their receptor genes in the spinal cord (**Supplementary Table 2**) were taken from the previously published studies (14,44). The *Avp, Avpr1b*, *Avpr2, Oxt* and *Pomc* genes were not included into statistical analysis because low expression levels of these genes in the spinal cord.

### Immunohistochemistry and estimation of the number of AVP neurons

#### Tissue preparation

Animals were deeply anaesthetized with pentobarbital and transcardially perfused with 150 mL of 0.1 M phosphate buffer (pH 7.4) for vascular rinse, followed by 250 mL of a fixative solution containing 4% paraformaldehyde in phosphate buffer. All rats were perfused between 2 and 5 p.m. Brains were removed from the skulls, immersed in fixative for 2 hours and infiltrated in 10% sucrose solution for 36 hours at 4°C. After removal of the frontal and occipital poles, the remaining tissue blocks were mounted on a vibratome and sectioned at 40 μm in the coronal plane. All sections cut through the hypothalamus posterior to the anterior commissure were collected using a systematic random sampling procedure (45). They were stored at -20°C in cryoprotectant (30% sucrose, 30% ethylene glycol, 0.25 mM polyvinylpyrrolidone in PBS) until use.

#### Immunohistochemistry and Nissl staining

From each brain, two sets of adjacent sections containing PVN were formed: (1) every second section was systematically sampled for immunostaining against AVP; (2) remaining sections were used for Nissl staining. In immunostaining procedure, the sections were washed three times in PBS and incubated in preheated (80°C) sodium citrate solution for 20 minutes for antigen retrieval. The endogenous peroxidases were removed by immersing the sections in 1% H_2_O_2_ solution for 15 minutes. After that the sections were washed six times in PBS (5 minutes each) and blocked for unspecific staining in PBS containing 10% normal goat serum (NGS) and 0.5% Triton X-100 for 2 hours at room temperature. The sections were then incubated with a rabbit polyclonal antibody against AVP (Merck Millipore AB1565, 1:4000 dilution in PBS) for 48 hours at 4°C. Subsequently, the sections were washed three times (10 minutes each) in 1%-NGS in PBS and then, incubated with a biotinylated horse anti-rabbit antibody (Vector Laboratories, BP-1100, 1:200 dilution) for 60 minutes. Sections were washed three times (10 minutes each) and incubated with avidin-biotin-peroxidase complex (1:400 dilution, Vector laboratories, Vectastain Elite ABC kit) for 45 minutes. After three times of washes the sections were recycled in secondary antibody and ABC for 45 and 30 minutes, respectively. The peroxidase reaction was obtained by incubation of sections in the solution of 3,3’-diaminobenzidine (DAB, 1 mg/ml) and H_2_O_2_ (0.08%) in PBS. Specificity of the immune reactions was controlled by omitting the incubation step with primary antisera. The sections were washed four times in PBS (10 minutes each) and then mounted on gelatin-coated slides and dried overnight. The sections were dehydrated in a series of ethanol solutions (50%, 70%, 96% and 100%), cleared with xylene and coverslipped using Histomount (National Diagnostics, Atlanta, GA, USA).

The second set of systematically sampled sections was stained for Nissl. The sections were mounted on gelatin-coated slides and air-dried. They were then stained with Giemsa dye (46), dehydrated in a series of ethanol solutions and coverslipped.

#### Estimation of the number of AVP-containing neurons

The AVP-stained sections containing the hypothalamic region were visualized using an Olympus BH-2 microscope equipped with a computer-controlled motorized stage and an MT-12 microcator (CAST grid system, Olympus Denmark). The boundaries of the PVN complex and its subdivisions were defined using the previously described cytoarchitectonic criteria (47,48) and the set of adjacent Nissl-stained sections. According to the terminology proposed in these studies, the magnocellular component of the PVN consists of three subdivisions, the anterior, medial, and posterior. In the present study, we analyzed only the posterior subdivision (PVNpm), one of the main sources of AVP afferents in the posterior pituitary. The number of AVP-stained neurons was estimated in all sampled sections in which the PVNpm was visualized (4-6 sections per animal) using the optical fractionator method (49). The region of interest was delineated using a 10× objective lens and cell counting was performed using a 100× oil-immersion lens. Starting at an arbitrary position, visual fields were systematically sampled with a step size of 100 µm along the x- and y-axes. The area of the counting frame was 1486 µm^2^. Starting 2 μm below the upper surface of the section (guard zone), cell counts were performed using a fixed height of 10 μm in all dissector frames. Neurons were counted separately in the left and right sides of the hypothalamus, as well as in the rostral and caudal halves of the PVNpm. Since there are no accepted cytoarchitectonic or anatomical criteria for delineating PVNpm subdivisions, cell counts from the anterior half of the sections were assigned to the rostral PVNpm and those from the posterior half were assigned to the caudal PVNpm. In addition, the total number of AVP cells in the left and right PVNpm was estimated for each animal.

### Immunohistochemistry of AVP-hM3Dq-mCherry transgenic rats

To investigate whether CNO activates AVP neurons, brains from the transgenic rats were used for immunofluorescent staining. The rats were first injected with either CNO (1 mg/mL/kg, i.p., n = 5) or vehicle (1 mL/kg, i.p., n = 5). Ninety minutes following the injection. the rats were deeply anaesthetized and transcardially perfused with 4% paraformaldehyde in 0.1M phosphate-buffered saline (PBS) and their brains were postfixed in the same solution for an overnight before being transferred to 0.1 M PBS containing 30% sucrose. The part of the brain that contains the hypothalamus was then coronally sectioned at 40 μm using a microtome (Microm HM450, ThermoScientific, Roskilde, Denmark). Every third consecutive sections were used for immunofluorescence to identify AVP and c-Fos expression. The sections were first incubated for 48 h at 4°C in one of the primary antibodies: guinea pig anti-Vasopressin (1:2000; Synaptic Systems, #403004), and mouse anti-c-Fos (1:500; Abcam, #ab208942), and then they were incubated in a secondary antibody for 1 h at room temperature. Since the AVP neurons in the AVP-hM3Dq-mCherry transgenic rat brains were pre-labeled red, the secondary antibodies used were tagged with green fluorophore, which was goat anti-guinea pig Alexa Flora 488 (1:500; Thermo Fisher Scientific, #A11073) or goat anti-mouse Alexa Flora 488 (1:500; Thermo Fisher Scientific, #A11029). Images were captured at 20X magnification using a DM600B Leica microscope (Leica Microsystems, Wetzlar, Germany). Positive cell numbers were counted using Image J cell counter plugin. The proportion of AVP-, and c-Fos-immunopositive neurons was calculated in relation to the Cherry-positive AVP neurons.

### Analysis of HL-PA

The HL-PA size and the side of the flexed limb were assessed before and 1 and 3 h after administration of neurohormones as described elsewhere (5,14,16). Briefly, the measurements were performed under isoflurane (1.5% isoflurane in a mixture of 65% nitrous oxide and 35% oxygen) anesthesia. The level of anesthesia was characterized by a barely perceptible corneal reflex and a lack of overall muscle tone. The anesthetized rat was placed in the prone position on the 1-mm grid paper.

In the hands-on analysis, the hip and knee joints were straightened by gently pulling the hindlimbs backwards for 1 cm to reach the same level. Then, the hindlimbs were set free and the magnitude of postural asymmetry was measured in millimeters as the length of the projection of the line connecting symmetric hindlimb distal points (digits 2-4) on the longitudinal axis of the rat. The procedure was repeated three times in immediate succession.

In the hands-off method, silk threads were glued to the nails of the middle three toes of each hindlimb, and their other ends were tied to one of two hooks attached to the movable platform that was operated by a micromanipulator constructed in the laboratory (14). To reduce potential friction between the hindlimbs and the surface with changes in their position during stretching and after releasing them, the bench under the rat was covered with plastic sheet and the movable platform was raised up to form a 10° angle between the threads and the bench surface. The limbs were adjusted to lie symmetrically and stretching was performed over a distance of 1.5 cm at a rate of 2 cm/sec. The threads then were relaxed, the limbs were released and the resulting HL-PA was photographed. The procedure was repeated three times in succession, and the HL-PA values for a given rat were used in statistical analyses.

The limb that projected over a shorter distance from the trunk was considered to be flexed. The HL-PA was measured in mm with negative and positive HL-PA values that were assigned to rats with the left and right hindlimb flexion, respectively. This measure, the postural asymmetry size (PAS) shows the HL-PA value and flexion side. The PAS cannot be used for analysis of rat groups with the similar number of left or right flexion. In the latter case the HL-PA value is about zero. Therefore, the HL-PA was also assessed by the magnitude of postural asymmetry (MPA) that shows absolute flexion size. The MPA does not show a flexion side. These two measures are obviously dependent; however, they are not redundant and for this reason, they are required for characterization of the HL-PA.

### Analysis of hindlimb resistance to stretch

Stretching force was analyzed before and 1 and 3 h after administration of peptide neurohormones using the micromanipulator-controlled force meter device constructed in the laboratory (5). Two Mark-10 digital force gauges (model M5-05, Mark-10 Corporation, USA) with a force resolution of 50 mg were fixed on a movable platform operated by a micromanipulator. Three 3-0 silk threads were glued to the nails of the middle three toes of each hindlimb, and their other ends were hooked to one of two force gauges. The flexed leg of the rat in the prone position was manually stretched to the level of the extended leg; this position was taken as 0 mm point. Then both hind limbs were stretched caudally, moving the platform by micromanipulator at a constant rate of 5 mm/sec for 10 mm. No, or very little, trunk movement was observed with stretching for the first 10 mm; therefore, the data recorded for this distance were included in statistical analysis. The forces (in grams) detected by each of the two gauges were simultaneously recorded (100 Hz frequency) during stretching. Three successive ramp-hold-return stretches were performed as technical replicates. Because the entire hindlimb was stretched, the measured resistance was characteristic of the passive musculo-articular resistance integrated for hindlimb joints and muscles (50–52). The resistance analyzed could have both neurogenic and mechanical components, but their respective contributions were not distinguished in the experimental design. The resistance was measured as the amount of mechanical work W_L_ and W_R_ to stretch the left- and right hindlimbs, where W was stretching force integrated over the stretching distance interval from 0 to 10 mm.

### Statistical Analysis

Experimental data were processed and statistically analyzed after completion of the experiments by the statisticians who were not involved in their execution. No intermediate assessment was performed to avoid any bias in data acquisition. The asymmetry data were obtained by unbiased hand-off method and unbiased registration of stretching force.

#### Molecular Analysis of Expression levels

The Lilliefors and Levene’s tests revealed deviations from normality and differences in the variances between the rat groups, respectively, for the expression levels and the asymmetry index (AI_L/R_ = log_2_L/R, where L and R were gene expression levels in the left and right halves of hypothalamus or spinal cord, respectively) of several genes in each rat group. The mRNA levels were compared separately for the left and right halves of the hypothalamus between i) the CNO-treated and vehicle-treated groups, and ii) UBI and sham surgery groups using Mann–Whitney test followed by Bonferroni correction for a number of tests.

The AI_L/R_ was computed for each gene and each individual animal. Median of the AI_L/R_ values represented the asymmetry at group-level. The AI_L/R_ was compared between the CNO- and vehicle-treated groups, and the UBI and sham surgery groups using Mann–Whitney test followed by a Bonferroni correction for multiple tests (n = 47-48 and 27 for the hypothalamus and spinal cord, respectively). Because no significant differences in the AI_L/R_ between the compared groups in both rat cohorts were revealed, the groups were combined respectively, and data for the pooled CNO-vehicle-treated group and the pooled UBI-sham surgery group were used for analysis of lateralization of gene expression using the AI_L/R_. Note that, in the “Gene Co-expression Patterns” section, correlations were computed for gene expression levels, but not for AI_L/R_. The rat groups were analyzed separately. One-sample version of non-parametric Wilcoxon signed-rank test was applied to compare the AI_L/R_ with zero followed by Bonferroni multiple testing correction. For the UBI-sham surgery cohort, we took data on the expression levels of 28 hypothalamic genes and 20 spinal cord genes from our previous studies (14,17,44).

#### Immunohistochemistry

Pairwise Pearson correlation coefficients were computed between the AVP-positive cell counts in left and right rostral and caudal subareas of the PVN (n = 21 rats). To compare two sets of dependent correlations—those involving the left rostral subarea versus those involving the right rostral and both caudal subareas—we applied Fisher’s R-to-Z transformation. The resulting Z-scores were then compared using a paired t-test, accounting for the non-independence of the correlation estimates. This statistical approach allowed for a valid comparison of correlation strength across functionally lateralized subregions.

#### Processing of physiological and molecular gene co-expression data

##### Bayesian framework

Predictors and outcomes were scaled and centered before we fitted Bayesian regression models via full Bayesian framework by calling *Stan 2.32.7* (Stan Development Team (2025). RStan: the R interface to Stan. R package version 2.32.7 https://mc-stan.org/) from *R 4.4.3* (R Core Team (2025). R: A language and environment for statistical computing. R Foundation for Statistical Computing, Vienna, Austria. URL https://www.R-project.org/) using the *brms 2.22.0* (53) interface. To reduce the influence of outliers, models used Student’s *t* response distribution with identity link function unless explicitly stated otherwise. Models had no intercepts with indexing approach to predictors (54). Default priors were provided by the *brms* according to Stan recommendations (Gelman A. 2019. Prior Choice Recommendations. Stan-dev/stan. https://github.com/stan-dev/stan/wiki/Prior-Choice-Recommendations. Accessed September 2022). Intercepts, residual SD and group-level SD were determined from the weakly informative prior student_t(3, 0, 3). Group-level effects were determined from the very weak informative prior normal(0, 2). Four MCMC chains of 40000 iterations were simulated for each model, with a warm-up of 20000 runs to ensure that effective sample size for each estimated parameter exceeded 10000 (55) producing stable estimates of 95% highest posterior density credible intervals (HPD). MCMC diagnostics were performed according to the Stan manual. P-values, adjusted using the multivariate t distribution with the same covariance structure as the estimates, were produced by frequentist summary in *emmeans 1.11.0* (Lenth R. 2025. emmeans: Estimated Marginal Means, aka Least-Squares Means. R package version 1.11.0. https://cran.r-project.org/package=emmeans. Accessed in 2025) together with the medians of the posterior distribution and 95% HPD. The asymmetry and contrast between groups were defined as significant if the corresponding 95% HPD did not include zero and the adjusted P-value was ≤ 0.05.

Stretching-force signals from the left and right hind limbs were smoothed with a loess estimator (span = 0.4, family = “symmetric”). Mechanical work for left limb (W_L_) and right limb (W_R_) was calculated by integrating the smoothed curve over a stretch distance of 0–10 mm. Asymmetry was expressed as the work difference W_L-R_ = W_L_ − W_R_.

Doses of GnRH and CCK-8 were analysed as a covariate. Across the readout models, the estimated log-dose slopes (median [95 % HPD]) for each peptide were effectively zero. All 95% HPD intervals overlapped zero, and pairwise contrasts between drugs also spanned zero. Therefore, within the tested dose range, peptide dose did not affect either metric analysed. Because log-dose added negligible explanatory power beyond the Time × Drug interaction—and to keep our models parsimonious—we omitted Dose.Log from the final analyses.

##### Gene co-expression patterns

For the hypothalamus and spinal cord separately, and for each cohort separately, genes were categorized into the LdN (median AI_L/R_ > 0) and RdN (median AI_L/R_ < 0).

The correlation structure (gene-gene co-expression pattern) for each area, side, between the sides, and across the areas was examined using the Spearman’s rank correlation coefficient calculated for all gene pairs. The pattern of interactions between genes was characterized by the coordination strength (magnitude of correlations or the absolute value of the correlation coefficient averaged across pairwise correlations) and the proportion of positive correlations.

Robust and unbiased P values were determined in the absence of distributional assumptions by permutation testing. A permutation procedure was employed to characterize the distribution of each statistical test under the null hypothesis of non-replication and non-preservation. Permutation test (Canty A, Ripley BD. 2022. boot: Bootstrap R (S-Plus) Functions. R package version 1.3-28.1.) with R=10^6^ bootstrap replicates implemented in the R/boot package was used to analyze the data. To generate null distribution, we permuted the data across i) *rat identification numbers (IDs)*, ii) *Treatment (UBI and sham surgery; CNO and vehicle)*; iii) *Module* (left and right) within each individual rat; and iv) *area* (hypothalamus and spinal cord).

The R/boot.pval package was used to compute the P-value (56). P-values were adjusted using Benjamini-Hochberg family-wise multiple test correction that was applied separately for all tasks.

### Data and code availability

Data supporting the findings of this study and all codes used for analysis are available within the article, its Supporting Information and on https://github.com/YaromirKo/biostatistics-GnRH_TRH_CCK.

## RESULTS

### Lateralized gene expression in the hypothalamus

Gene expression asymmetry was quantified using the asymmetry index AI_L/R_ = log₂(L/R), where L and R represent expression levels on the left and right hypothalamus, respectively (for details, see “**Materials and Methods**”). The analyzed set consisted of 48 genes including most hypothalamic neurohormones and neuropeptides genes, their receptor genes and neuroplasticity-related genes (**Supplementary Table 1**). Neuroplasticity-related genes—transcriptional regulators, glutamate system components, and genes involved in synaptic remodeling, axon growth, and neuroimmune signaling (for details of gene selection, see “**Materials and Methods**”)—were included to assess their role in neurohormonal lateralization.

The AI_L/R_ was computed for each gene and each individual animal. Rat cohorts analyzed were: i) naïve Wistar rats (n = 11); ii) transgenic AVP-hM3Dq-mCherry rats, which were treated for 3 h with either clozapine-N-oxide (CNO; n = 11) to stimulate transgene or vehicle (n = 11); and iii) rats with complete thoracic spinal cord transection followed by UBI (n = 12) or sham-surgery (n = 11) (**Supplementary Figures 1 and 2**).

We previously demonstrated that nearly all mCherry-positive neurons were co-labeled with AVP antibodies in the magnocellular division of the PVN, as well as in the supraoptic nucleus and suprachiasmatic nucleus (20). Approximately 90% of mCherry-expressing neurons in these nuclei showed c-Fos expression after CNO treatment compared to 5–17% in control animals, and chemogenetic stimulation induced AVP release (20). Consistent with this study, 94.0 ± 3.7% and 92.7 ± 5.6% of mCherry-positive neurons co-expressed AVP in the PVN of the vehicle- and CNO-treated groups, respectively (shown as mean ± standard deviation). The same hold true for supraoptic nuclei (97.0 ± 3.6% and 96.2 ± 2.2%, respectively). CNO administration did not alter mCherry expression levels.

Neuronal activation was assessed by c-Fos immunoreactivity. Ninety minutes after CNO injection, robust induction of c-Fos was observed in the PVN and supraoptic nuclei compared to vehicle controls. Specifically, 80.3 ± 7.5% of mCherry-positive AVP neurons in the PVN and 81.4 ± 9.0% in the SON expressed c-Fos, versus 9.6 ± 4.9% and 5.7 ± 5.2%, respectively, in the control group (shown as mean ± standard deviation). Differences between the groups were highly significant (P < 0.001, unpaired two-tailed t-test), confirming successful activation of AVP neurons via hM3Dq.

Left UBI was performed because its stronger effects *vs*. those of right UBI on gene expression (14,15). Samples were collected at the time point that was five hours after spinal transection and three hours after UBI or sham surgery.

The AI_L/R_ values did not significantly differ between sexes or between CNO and vehicle treated transgenic rats, nor between UBI and sham groups (Mann–Whitney test followed by Bonferroni correction). Two subgroups of each cohort were therefore pooled (the transgenic cohort: n = 22; the UBI-sham cohort: n = 23) for the statistical analysis. The AI_L/R_ values for individual genes in three cohorts are presented in **Supplementary Figure 2A-C**. Of the 47 genes analyzed, 21 showed consistent directionality of asymmetry across all three cohorts, and 37 were consistent between Wistar and transgenic cohorts. AI_L/R_ values significantly correlated between Wistar and transgenic animals (Pearson *R* = 0.52, P = 2e-04; Spearman ρ = 0.53, P = 1.2e-04) (**Supplementary Figure 2D**).

Using two stringent criteria—significant AI_L/R_ (a two-sided Wilcoxon signed rank exact test: Bonferroni-adjusted P < 0.05) in the combined cohort (n = 56 rats), and the same AI_L/R_ sign across all cohorts—we identified 11 significantly lateralized genes (**Table 1**). The latter criterion was imposed to rule out the predominance of any single cohort over others in the pooled sample. Expression of *Pomc*, *Trh* and *Oprk1* was lateralized to the left, whereas that of the *Cck*, *Mas1*, *Pnoc*, *Gnrh1*, *Ghrl*, *Ren*, *Nfkbia* and *Agtrap* to the right. *Pomc*, *Mas1*, and *Cck* exhibited approximately 2-fold expression differences between hemispheres.

**Table 1.**
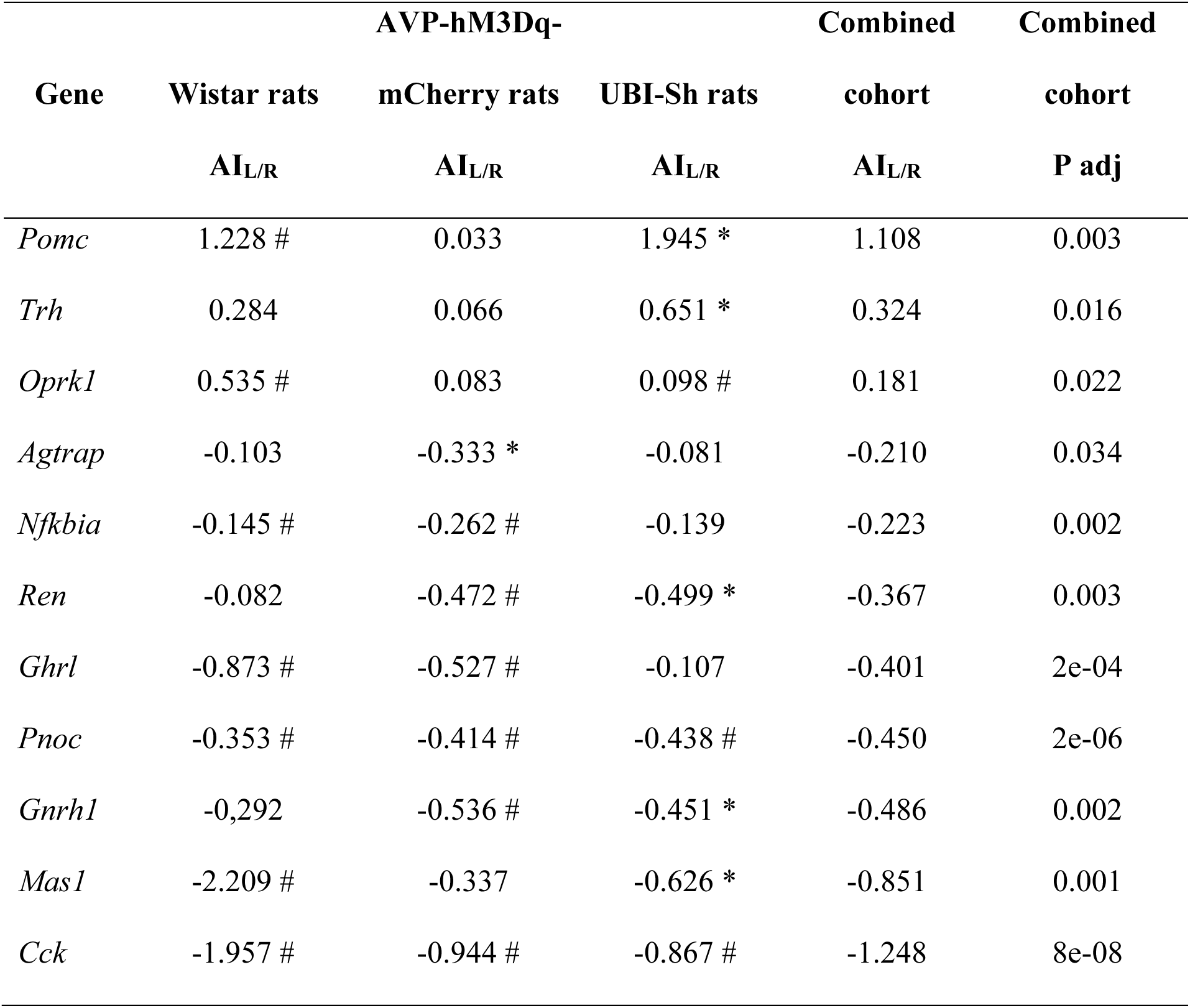
Lateralized expression of the neurohormonal and plasticity-related genes in the rat hypothalamus. Shown is the asymmetry index (AI_L/R_ = log_2_[L/R], where L and R are the median expression levels in the left and right hypothalamus) for the naïve Wistar cohort (n = 11), the combined transgenic AVP-hM3Dq-mCherry cohort of rats treated with CNO or vehicle, and the combined UBI - sham surgery cohort. In transgenic cohort, there were no differences in the AI_L/R_ between female and male subgroups, and between CNO (n = 11) and vehicle (n = 11) treated rats in the AI_L/R_; therefore, they were combined (n = 22) for statistical analysis of the asymmetry of expression of individual genes (Mann–Whitney test followed by a Bonferroni correction). In the UBI and sham surgery cohort, **t**here were no differences in the AI_L/R_ between UBI (n = 12) and sham surgery (n = 11) groups; therefore, they were combined (n = 23) for statistical analysis of the asymmetry of expression of individual genes (Mann–Whitney test followed by a Bonferroni correction). Data for these three cohorts were also pooled together and analyzed as the combined cohort (n = 56 rats). * and #, significant (P < 0.05 after Bonferroni correction) and nominally significant lateralization in a given cohort. Data are shown for eleven genes that were significantly lateralized, out of 47 genes analyzed. Lateralization was considered to be significant if P adjusted (P adj) by Bonferroni correction was < 0.05 in the combined cohort, and the direction of lateralization was the same in three experimental cohorts. The P-values were computed for 47 genes using a two-sided Wilcoxon signed rank exact test and adjusted using the Bonferroni correction in three cohorts separately and in the combined cohort.

In transgenic rats, no significant changes in expression of individual genes were observed after correction between CNO and vehicle groups in either hemisphere. However, the proportion of up- and downregulated genes differed significantly between the sides after CNO treatment. In the left hypothalamus, 33 genes had lower and 14 had higher median expression in the CNO group compared to the vehicle group. In contrast, in the right hypothalamus, only 13 genes had lower expression whereas 34 genes had higher expression levels in the CNO group compared to the vehicle group. This asymmetry was statistically significant (Fisher’s exact test: P = 7e-5), suggesting preferential gene downregulation in the left and upregulation in the right hypothalamus following stimulation of AVP neurons.

For UBI effects, no individual genes passed the significance threshold after correction. Still, a significant majority of genes were downregulated on both sides, with more pronounced effects on the left: 35 out of 48 genes showed decreased expression on the left side (Fisher p = 0.035), and 37 out of 48 genes on the right side (P = 0.010). Furthermore, 15 genes showed nominal significance on the left vs. only 2 on the right (P = 9e-4), indicating stronger overall effects of left UBI on ipsilateral hypothalamic gene expression.

### Hypothalamic gene co-expression networks

To explore how gene expression is coordinated within the left and right hypothalamus, we analyzed co-expression networks stratified by lateralization. Genes were grouped by the sign of their AI_L/R_: genes with AI_L/R_ > 0 were classified as left-dominant (LdN), while those with AI_L/R_ < 0 were classified as right-dominant (RdN) (**Supplementary Figure 2**). Genes were further grouped by hemisphere of expression into the left module (LM) and right module (RM), each containing both LdN and RdN genes.

For each module, we computed pairwise Spearman correlations across all gene pairs. Two key features were derived: (1) **coordination strength**, defined as the average absolute correlation coefficient among gene pairs, and (2) **proportion of positive correlations**, reflecting the directionality of network interactions. Robust P values were estimated via permutation testing to account for non-normal distributions.

We compared intra-network (LdN–LdN and RdN–RdN) and inter-network (LdN–RdN) co-expression within each hemisphere (LM and RM) and across treatment groups (**Figures 1-3**). These analyses were performed separately for all rat cohorts and their subgroups: the Wistar cohort, transgenic rats (vehicle vs. CNO), and UBI vs. sham-lesioned groups. Notably, correlation metrics were computed from raw expression values, not AI_L/R_, which showed no subgroup differences in either cohort.

**Figure 1.**
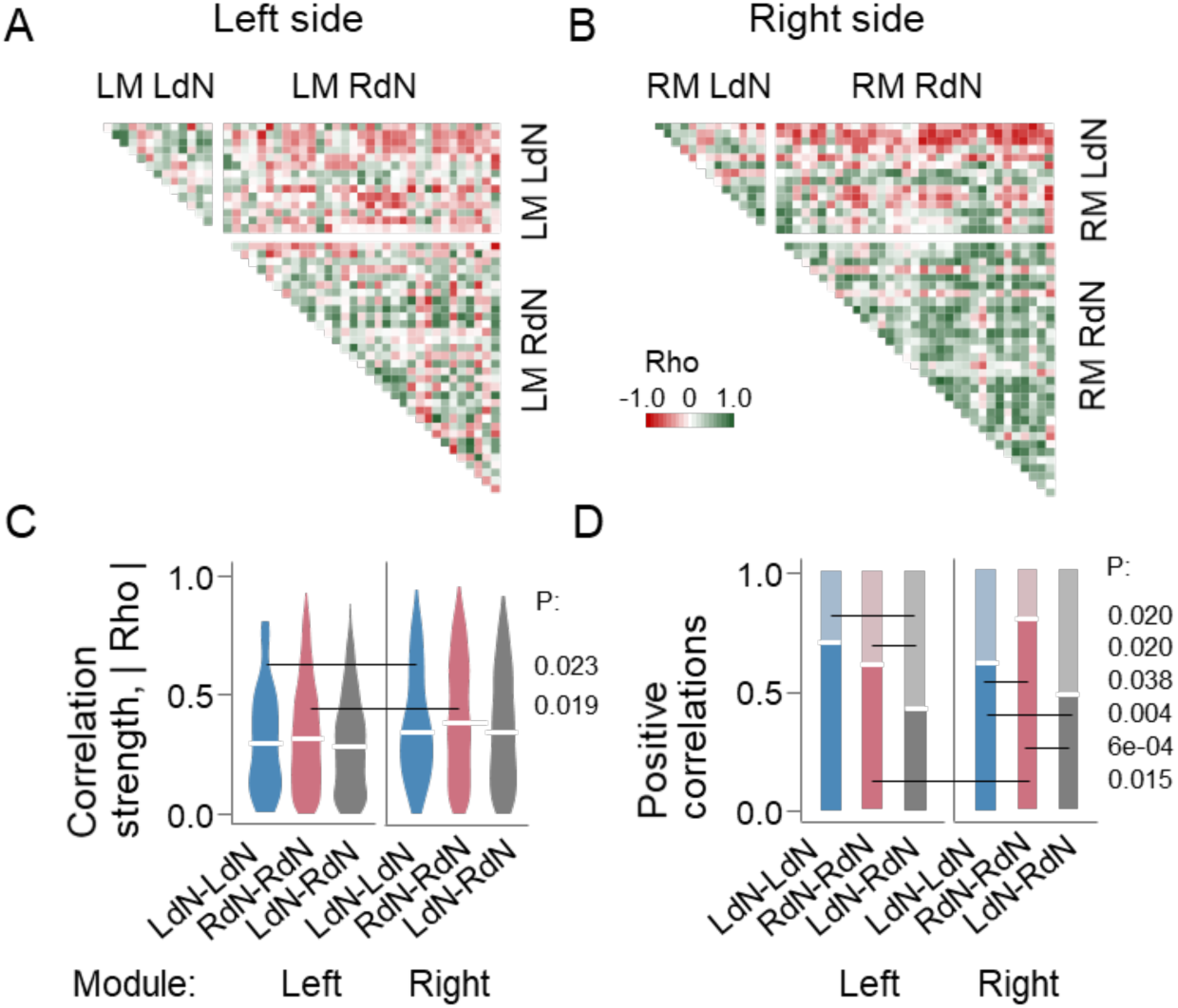
Gene co-expression patterns in the hypothalamus of naïve Wistar rats (n = 11). (**A**,**B**) Heatmaps for Spearman’s rank coefficients for pairwise gene-gene correlations in the left (LM) and right (RM) modules. (**C,D**) Patterns of intra-modular correlations internal for each the LdN (LdN-LdN) and RdN (RdN-RdN), and those between the networks (LdN-RdN) for the LM and RM. In (**C**), strengths of pairwise correlations (correlation coefficient |Rho|) are presented as violin plots with white line indicating the coordination strength (the absolute Rho value averaged across correlations). In (**D**), the proportion of positive correlations shown as white lines. The correlation patterns were compared within each LM and RM and each pattern was compared between the modules. Shown are P values that were determined by permutation testing with Benjamini-Hochberg family-wise multiple test correction.

Two co-expression features general for all cohorts were revealed. First, coordination strength was higher in the right hypothalamus compared to the left at most comparisons. This was statistically significant in the Wistar cohort (for LdN–LdN and RdN–RdN) (**Figure 1**), the transgenic CNO group (for RdN–RdN) (**Figure 2**), and both UBI and sham groups (for LdN–LdN) (**Figure 3**). Second, the proportion of positive correlations was higher for intra-network patterns (LdN–LdN and RdN–RdN) compared to the inter-network LdN–RdN pattern in 16 of 17 significant cases (Fisher’s exact test, two-tailed: P = 0.007). Significant differences were detected in LdN–LdN vs. LdN–RdN and RdN–RdN vs. LdN–RdN in both hemispheres of the Wistar cohort and both UBI and sham groups, RdN–RdN vs. LdN–RdN in both hemispheres of the vehicle group and in the right hemisphere of the CNO group, and LdN–LdN vs. LdN–RdN in the left hemisphere of the CNO group.

**Figure 2.**
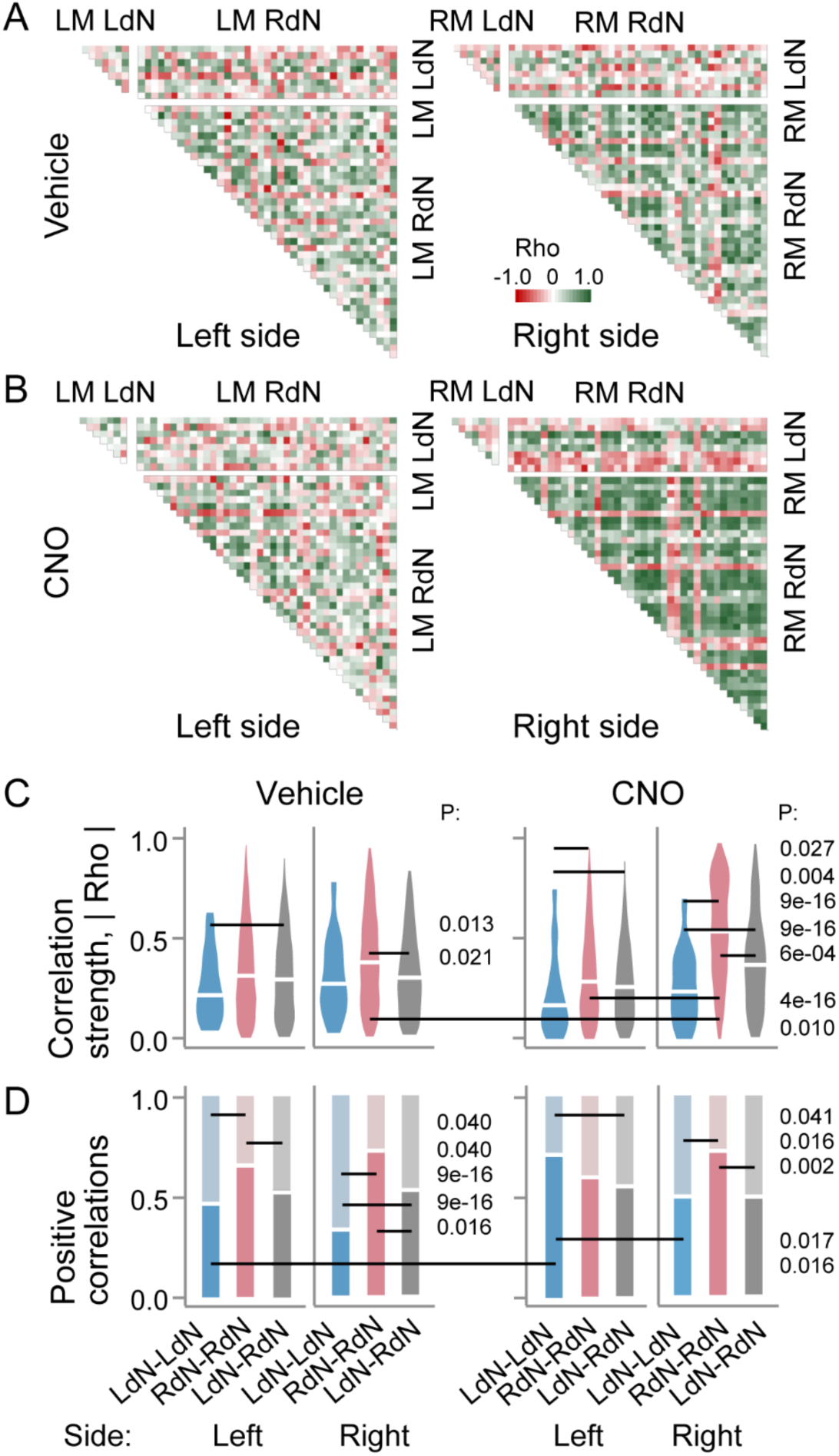
Gene co-expression patterns in the hypothalamus of the AVP-hM3Dq-mCherry rats treated with CNO (n = 11) or vehicle (n = 11) for 3. **h.** (**A,B**) Heatmaps for Spearman’s rank coefficients for pairwise gene-gene correlations in the LM and RM) of vehicle- and CNO-treated groups. (**C,D**) Patterns of intra-modular correlations internal for each the LdN (LdN-LdN) and RdN (RdN-RdN), and between the networks (LdN-RdN) are shown for the left and right modules of sham surgery and left UBI groups. In **C**, strengths of pairwise correlations (correlation coefficient |Rho|) are presented as violin plots with white line indicating the coordination strength (the absolute Rho value averaged across correlations). In **D**, the proportion of positive correlations shown as white lines. The correlation patterns were compared within each LM and RM and each pattern was compared between the modules, and between the vehicle- and CNO-treated rat groups. Shown are P values that were determined by permutation testing with Benjamini-Hochberg family-wise multiple test correction.

The conservation of these two correlation features across cohorts supports the existence of lateralized gene co-expression networks in the hypothalamus. These patterns are clearly visualized in the correlation heatmaps (**Figures 1-3**).

**Figure 3.**
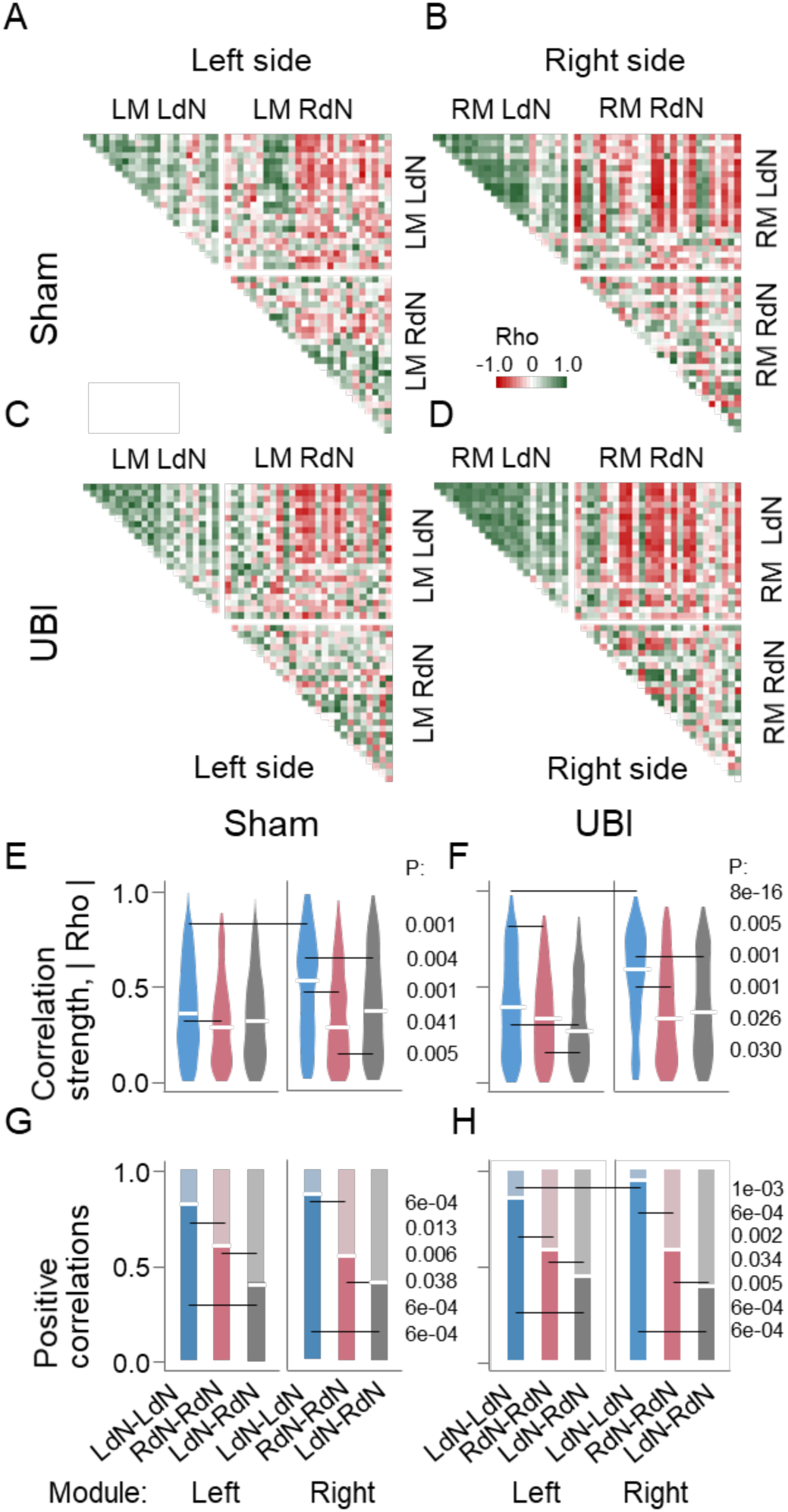
Gene co-expression patterns in the hypothalamus of sham surgery (n = 11) and UBI (n = 12) rats. Samples were collected five hours after spinal transection, which was three hours after UBI or sham surgery. (**A-D**) Heatmaps for Spearman’s rank coefficients for pairwise gene-gene correlations in the LM and RM of sham surgery and UBI groups. (**C,D**) Patterns of intra-modular correlations internal for each the LdN (LdN-LdN) and RdN (RdN-RdN), and between the networks (LdN-RdN) are shown for the LM and RM of the left sham surgery group and the left UBI group. In **E** and **F**, strengths of pairwise correlations (correlation coefficient |Rho|) are presented as violin plots with white line indicating the coordination strength (the absolute Rho value averaged across correlations). In **G** and **H**, the proportion of positive correlations shown as white lines. The correlation patterns were compared within each LM and RM and each pattern was compared between the modules, and between the sham surgery and UBI groups. Shown are P values that were determined by permutation testing with Benjamini-Hochberg family-wise multiple test correction.

Chemogenetic stimulation of AVP neurons produced robust asymmetric effects on network coordination (**Figure 2**). Compared to vehicle-treated controls, CNO-treated rats showed: i) significantly higher coordination strength in the RdN–RdN pattern in the right hypothalamus, and ii) a significantly greater proportion of positive correlations in the LdN–LdN pattern in the left hypothalamus. Gene co-expression patterns were practically symmetric in vehicle-treated rats – no significant left-right differences revealed. However, stimulation of AVP neurons resulted in a strong statistically significant increase in RdN–RdN coordination strength in the right hypothalamus compared to the left. No significant differences were revealed between male and female subgroups in the transgenic cohort (12 males vs. 10 females).

### Asymmetric rostro-caudal distribution of AVP neurons in the hypothalamus

AVP mediates the contralateral impact of left-sided brain injury via humoral pathways by producing asymmetric effects—flexion of the right hindlimb (14). Encoding of hemisphere-specific signals into AVP-mediated responses may depend on a lateralized organization of AVP-expressing neurons in the hypothalamus. However, gene expression analysis of bulk hypothalamic tissue samples revealed no significant asymmetry in AVP mRNA levels. To address this discrepancy, we compared the number of AVP-expressing neurons between the left and right PVN—a hypothalamic region known to be functionally organized along its rostro-caudal axis (47,57), using stereology-based quantitative immunohistochemistry. We focused on the posterior magnocellular subdivision of the PVN—PVNpm, one of the major sources of AVP projections to the posterior pituitary (58,59), and quantified AVP-immunopositive neurons in the left and right PVNpm across 21 rats.

Total AVP-positive neuron counts did not differ significantly between the left and right PVNpm (**Supplementary Table 3**). The asymmetry index AI also did not deviate from zero (one-sample *t*-test and Mann–Whitney test). Separate analysis of the rostral (anteroventromedial) and caudal (posterodorsolateral) PVNpm subdivisions (**Figure 4**) also did not reveal significant left-right differences in these parameters in either subregion, and the AI between the two regions. However, the non-directional asymmetry—represented by absolute AI values |AI|—was significantly greater in the rostral PVNpm compared to the caudal region (P = 0.044; *t*-test and Mann–Whitney test; **Figure 4B**). Thus, while total AVP neuron distribution is symmetric in the PVNpm, rostral and caudal subregions may differ in the degree of non-directional asymmetry, suggesting a lack of coordination between these compartments.

**Figure 4.**
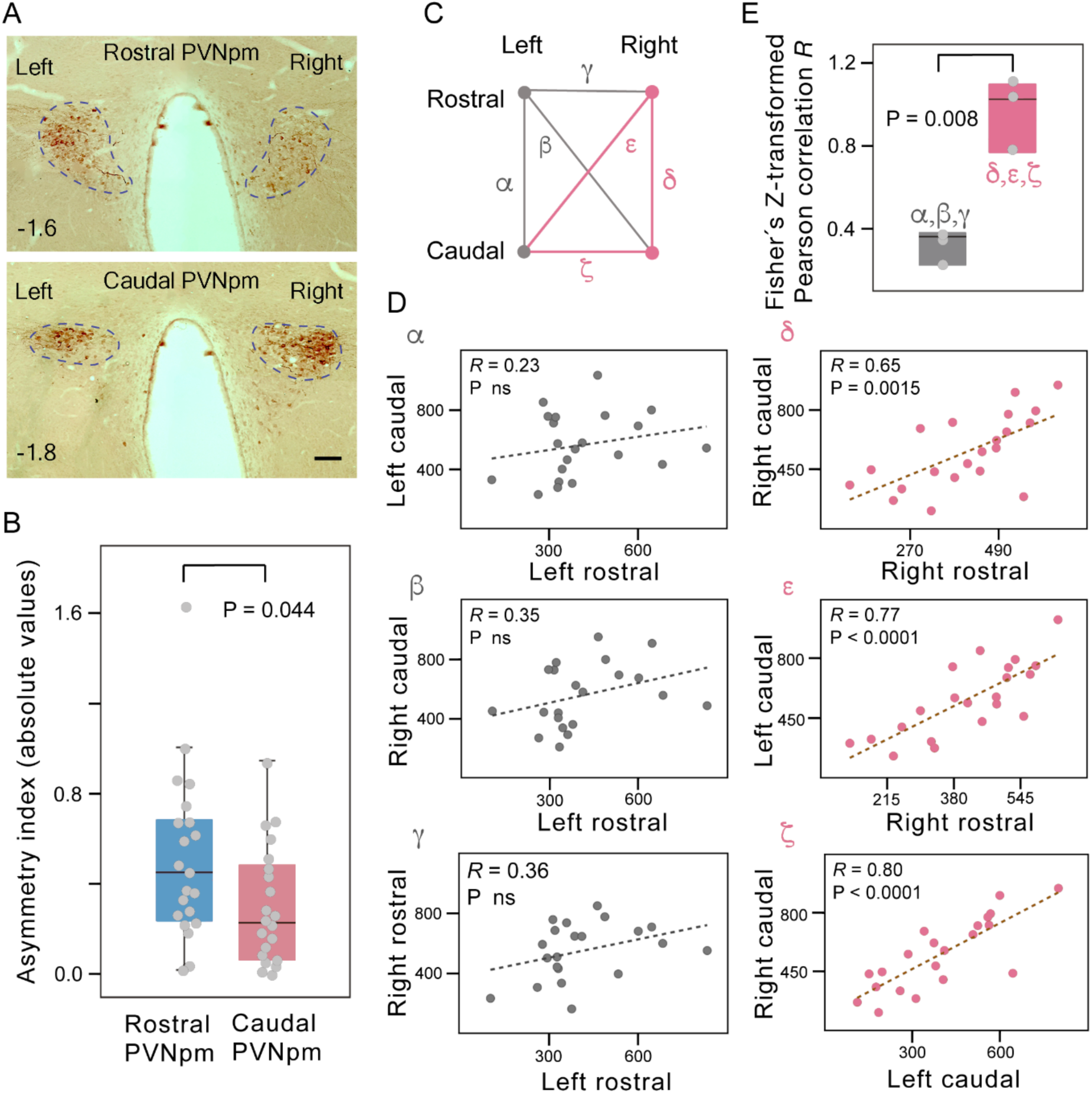
Lateralized coordination between the rostral and caudal groups of AVP-expressing cells in posterior magnocellular paraventricular nucleus (PVNpm). **(A)** Representative photographs of AVP-immunostained brain sections from a control rat taken from the rostral (top panel) and caudal (bottom panel) subareas of the PVNpm (delineated by a dashed line). The images show that the left-right distribution patterns differ between the rostral and caudal parts. Scale bar = 100 μm. The numbers on the left indicate distances (in mm) from bregma. **(B)** Comparison of non-directional asymmetry in AVP cell number between the rostral and caudal PVNpm subareas. In statistical analysis, rats of four experimental groups (control, sham-UBI, left-UBI and right-UBI) were pooled together (final n = 21), as no significant differences were found between the groups for either morphometric measure. The absolute value of the AI_L/R_ for AVP-positive cell numbers |AI| was determined in the rostral and caudal parts of the PVNpm of each animal. It was significantly greater in the rostral PVNpm compared to its caudal part (P = 0.044; t-test and Mann–Whitney test). **(C)** Schematic representation of a coordination network formed by the four magnocellular PVN domains with high AVP expression: the left and right rostral subareas and the left and right caudal subareas. Two sets of links in this network (Set A – α, β, γ, centered on the left rostral PVNpm, and Set B – 8, χ, β, centered on the right rostral PVNpm) can be distinguished on the basis of the data shown in **(B)** and **(D)**. **(D)** Correlations in the number of AVP cells across four PVNpm subregions depicted in **(C)**. Correlation analyses revealed that all three links in the Set A are weak and nonsignificant, whereas the links in the Set B are strong and highly significant (Pearson and Spearman correlations yielded similar results). **(E)** Comparison of correlation strengths between the two subnetworks. Fisher Z-transformed Pearson correlation coefficients were computed for Set A (panels α, β, γ in **(D)**) and Set B (panels 8, χ, β in **(D)**). A paired t-test revealed a significant difference between the sets (t(2) = 11.35, p = 0.008), indicating stronger coordination among the PVNpm subareas constituting the Set B than among those constituting the Set A.

Following correlation analysis of AVP neuron numbers between subregions and hemispheres revealed a distinct organizational pattern (**Figure 4C-E**). Correlations involving the left rostral PVNpm were weak and nonsignificant (denoted as α, β, γ). In contrast, strong and significant correlations were found among the right rostral PVNpm and both caudal subregions, and between the left and right caudal PVNpm (denoted as δ, ε, ζ). The mean correlation coefficient among the δ, ε, and ζ pairs was approximately threefold higher than that of the α, β, γ group (paired *t*-test on Fisher Z-transformed *R* values; P = 0.008; **Figure 4E**). These findings suggest that AVP neuron numbers are coordinated between the right rostral and caudal subregions, whereas the left rostral PVNpm is more autonomic. This lateralized connectivity along the rostro-caudal PVN axis suggestsdistinct functional specializations of the left and right rostral and caudal PVNpm subdivisions in side-specific neuroendocrine signaling.

### Gene co-expression patterns across the hypothalamus and spinal cord

Side-specific hormonal interactions between the hypothalamus and peripheral endocrine targets may be reflected in coordinated gene co-expression patterns between the left and right sides of these areas and tissues. Previous study with a limited gene set suggested gene co-expression patterns across the hypothalamus and spinal cord that are formed through humoral pathway and affected by left UBI (17). In the present work, using an expanded dataset—including (i) genes encoding most hypothalamic neurohormones, and (ii) genes for the majority of spinal neuropeptides and their receptors—we validated side-specific hypothalamic–spinal cord coordination (**Supplementary Tables 1** and **3**). Data obtained from the UBI-sham surgery rats whose spinal cord were completely transected, were analyzed. The spinal cord LdN and RdN were defined for 27 neurohormonal, neuropeptide and plasticity-related genes on the basis of their AI (**Supplementary Figure 3**). We first computed ipsilateral Spearman’s correlations separately for LdN and RdN between the hypothalamic and spinal modules: LM–LM (left hypothalamus and left spinal cord), and RM–RM (right hypothalamus and right spinal cord) (**Supplementary Figure 4**). In sham-operated rats, striking asymmetries were observed in the proportion of positive correlations (**Supplementary Figure 4B,C**). For the LdN, the proportion was significantly higher on the left than the right side. Conversely, for the RdN, the proportion was higher on the right than on the left. Within each spinal module, the LdN pattern exceeded the RdN pattern in the LM, and the RdN exceeded the LdN in the RM. The LdN–LM and RdN–RM patterns in the proportion of positive correlations were comparable, as were the RdN–LM and LdN–RM patterns. Additionally, coordination strength was greater for LdN than for RdN in the RM pattern.

Left-sided UBI disrupted these hypothalamic–spinal interactions for only the LdN patterns but not the RdN patterns (**Supplementary Figure 4B,D**). UBI significantly reduced coordination strength for LdN in both LM and RM (i.e., on both sides), decreased the proportion of positive correlations in the left (ipsilesional) LdN pattern, while increasing the proportion on the right (contralesional) side.

By contrast, significant contralateral (diagonal) correlations—i.e., between the left hypothalamus and right spinal cord, and vice versa—were much fewer in number than ipsilateral correlations for both LdN and RdN (**Supplementary Figure 5**). These diagonal correlations were largely symmetric across both coordination strength and the proportion of positive correlations, and showed no significant differences between the α- and γ-patterns or the β- and δ-patterns depicted in **Supplementary Figure 5**, in either the sham or UBI groups.

In summary, gene co-expression patterns between the hypothalamus and spinal cord were strongly ipsilateral, lateralized, and distinct for LdN and RdN networks. These patterns were significantly perturbed by UBI in rats with fully transected spinal cords—indicating that coordination is mediated through humoral, not neural, pathways. The results confirm and expand upon our prior findings (14,17), supporting the existence of a functional, endocrine-mediated, left–right side-specific communication axis between the hypothalamus and spinal cord.

### Side-specific neurohormonal effects mediated via humoral pathway

We next investigated whether hypothalamic neurohormones—constituents lateralized gene co-expression networks—can invoke asymmetric signaling from the brain to peripheral endocrine targets mediated via the humoral pathway. We analyzed peripheral responses to three neurohormones—GnRH, CCK-8, and TRH—after their administration into the cisterna magna (**Figure 5**, **Supplementary Figure 6**). The HL-PA was used as a binary readout model that captures side-specific peripheral changes – asymmetry in hindlimb functions and posture, and that was previously validated in analysis of lateralized actions of neurohormones including opioid peptides and AVP (14,15,17). To isolate humoral signaling and exclude transmission of neurohormonal effects through descending neural pathways, the spinal cords were completely transected at the thoracic level prior to peptide administration. Asymmetry was assessed before, and 1 and 3 hours after injection, by measuring (i) passive musculo-articular resistance to hindlimb stretch (**Figure 5C**), and (ii) hindlimb postural asymmetry (**Supplementary Figure 6**). GnRH was injected at doses 0.1, 1, and 50 ng/rat, CCK-8 – at doses 0.1, 1, 10, and 30 ng/rat, while TRH at dose 1 ng/rat. The number of rats used in analysis is given in **Supplementary Table 4.**

**Figure 5.**
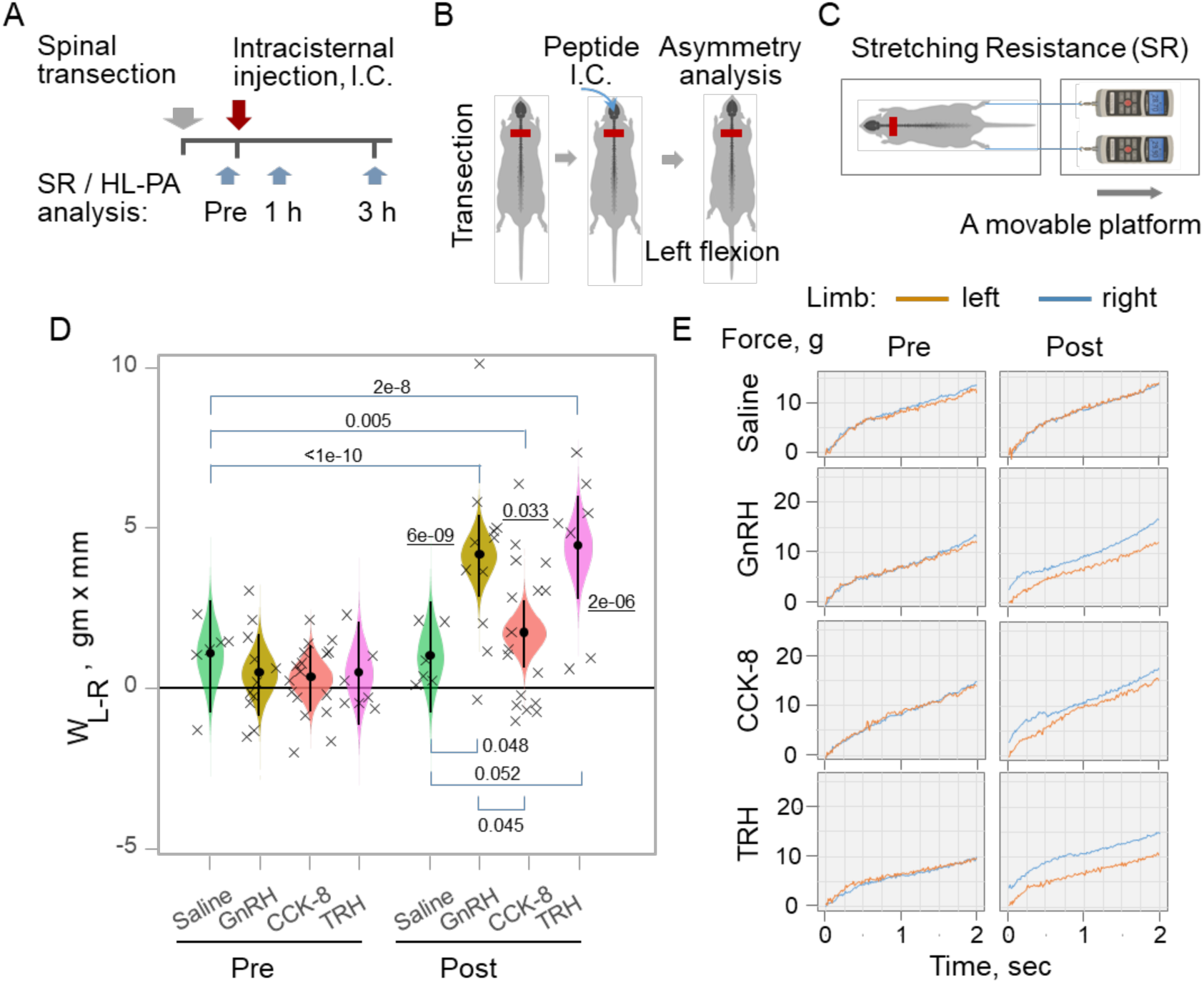
Asymmetry in hindlimb stretching resistance (SR) induced by neurohormones administered into the cisterna magna of rats with completely transected thoracic spinal cords. (**A**,**B**) Experimental design. The spinal cord was transected at the T2-T4 level, and then neurohormones or saline were administered intracisternally. Stretching resistance data are shown for asymmetry levels analyzed before (Pre) and 1 h after injection. The significance of the peptide effects was essentially the same for the 3 h post injection time point. The HL-PA data for these rats are shown in **Supplementary Figure 6**. (**C**) The stretching resistance was analyzed as the amount of mechanical work W required to stretch a hindlimb, calculated as the integral of stretching force over a distance of 0-10 mm. The force was measured using a micromanipulator-controlled force meter consisting of two digital force gauges fixed on a movable platform. (**D**) Differences in stretching force between the left and right hindlimbs W_L-R_ in gm ξ mm. The median (represented as black circles), 95% HPD (black lines), and posterior density (colored distribution) from Bayesian regression are used to plot W_L-R_. W_L-R_ values for individual rats are indicated by crosses, Asymmetry and contrasts among the groups were deemed significant, with a 95% HPD not encompassing zero and adjusted P values of ≤ 0.05. Adjusted P values are presented numerically on the plots for significant W_L-R_ asymmetries (P values are underlined), and for significant contrasts (pointed by brackets) in W_L-R_ between i) Pre and Post (1 h) measurements in the same group of animals; ii) groups of animals injected with peptides and saline; and iii) groups of animals injected with different peptides. Rats received saline (n = 6); GnRH at doses 0.1 (n = 3 rats), 1 (n = 6 rats), and 50 (n = 2 rats) ng/rat; CCK-8 at doses 0.1 (n = 2 rats), 1 (n = 6 rats), 10 (n = 2 rats), and 30 (n = 7 rats) ng/rat; or TRH at dose 1 ng/rat (n = 7 rats). No significant differences between the doses for GnRH were revealed and the data were combined (dose was included as covariate in statistical analysis; see Materials and Methods). The same holds true for CCK-8. (**E**) Representative traces of the stretching force recorded from the left and right hindlimbs before and after peptide administration.

### Hindlimb stretching resistance

Lateralized resistance to hindlimb stretch was quantified as the difference in mechanical work between the left and right hindlimbs as W_L-R_ = (W_L_ - W_R_), where W_L_ and W_R_ were the work applied to stretch the left and right hindlimbs, respectively. **Figure 5** shows data for the 1 h post injection time point; the peptide effects and their significance were essentially the same for the 3 h time point. **Figure 5E** shows representative force traces recorded before and after peptide administration. The force required to stretch each limb increased progressively with the degree of stretch. **Figure 5D** presents the median of each group, the 95% highest posterior density credible interval (HPD), and the corresponding posterior distribution, derived from Bayesian regression (see **Glossary**). The 95% HPD represents the interval within which the true group median is expected to fall with 95% probability—analogous to a traditional confidence interval. Animals were defined as significantly asymmetric if the 95% HPD did not include zero and the adjusted P-value for the relevant contrast was < 0.05. No significant asymmetry in hindlimb resistance was observed before injection or after saline treatment (**Figure 5D,E**). In contrast, significant left–right asymmetry was observed after administration of GnRH (P = 6e-9), CCK-8 (P = 0.033), and TRH (P = 2e-6), with the left hindlimb exhibiting greater resistance to stretch. There were no significant differences between the doses of GnRH, so the data were pooled for final analysis (see statistical section in Materials and Methods). The same holds true for CCK-8.

The contrasts in W_L-R_ were statistically significant both (i) within groups: comparing values before vs. after injection for all three neurohormones; and (ii) between groups: comparing peptide-treated animals to those receiving saline (**Figure 5D**). P-values (adjusted for multiple comparisons) reflect differences in median values between groups. For example, P = 0.048 for the contrast “Post: GnRH – Post: Saline” refers to the difference between the post-injection medians for GnRH and saline groups.

### HL-PA

HL-PA was analyzed using both the hands-on and hands-off methods of hindlimb stretching followed by photographic and/or visual recording of asymmetry in anesthetized animals (**Supplementary Figure 6**, **Supplementary Table 4**) (14,15,17). Data from these two methods correlated well with each other (17) and they produced essentially the same results; those presented in **Supplementary Figure 6** are for the hands-off assay. HL-PA was characterized by i) the magnitude of postural asymmetry (MPA) in mm, and ii) the size of postural asymmetry (PAS) in mm. In contrast to the MPA, the PAS shows the direction of the asymmetry; negative and positive PAS values are assigned to left and right hindlimb flexion, respectively.

Following central administration of GnRH, CCK-8, and TRH, a consistent HL-PA emerged, with the left hindlimb flexion (**Supplementary Figure 6B**,**C**). This shift was evident both 1 hour and 3 hours post-injection and mirrored the pattern observed in stretch resistance. The directionality of the response—favoring the left hindlimb—was consistent across peptides (**Supplementary Figure 6C**). The contrasts in the MPA and PAS for the three peptides were statistically significant both (i) within groups: comparing values before vs. after peptide injection; and (ii) between groups: comparing peptide-treated animals to those receiving saline (**Supplementary Figure 6D-G**).

The stretching resistance and HL-PA findings indicate that centrally administered neurohormones can produce asymmetric side-specific peripheral effects in the absence of spinal connectivity, implicating a humoral pathway.

## DISCUSSION

### Asymmetry of the Neuroendocrine Hypothalamus

Our findings support the hypothesis that the hypothalamus exhibits left–right asymmetry in the distribution and co-expression patterns of neurohormonal genes, which may underlie the encoding of hemisphere-specific neural signals into lateralized humoral outputs (**Figure 6**). Early studies have demonstrated functional hypothalamic asymmetry, particularly in the regulation of reproductive function and behavior (60–77). In neonatal female rats, exposure of the left or right hypothalamus to estrogen led to contrasting effects: defeminization or masculinization, respectively (68). In adult animals, unilateral hypothalamic lesions, hemigonadectomy, and pharmacological interventions revealed ipsilateral control of gonadotropin secretion and ovarian function, with a consistent right-sided dominance (60–67). A key mediator in these processes was GnRH, which has been shown to be lateralized to the right hypothalamus (69–77). Our finding of asymmetric Gnrh1 gene expression is consistent with these earlier observations.

**Figure 6.**
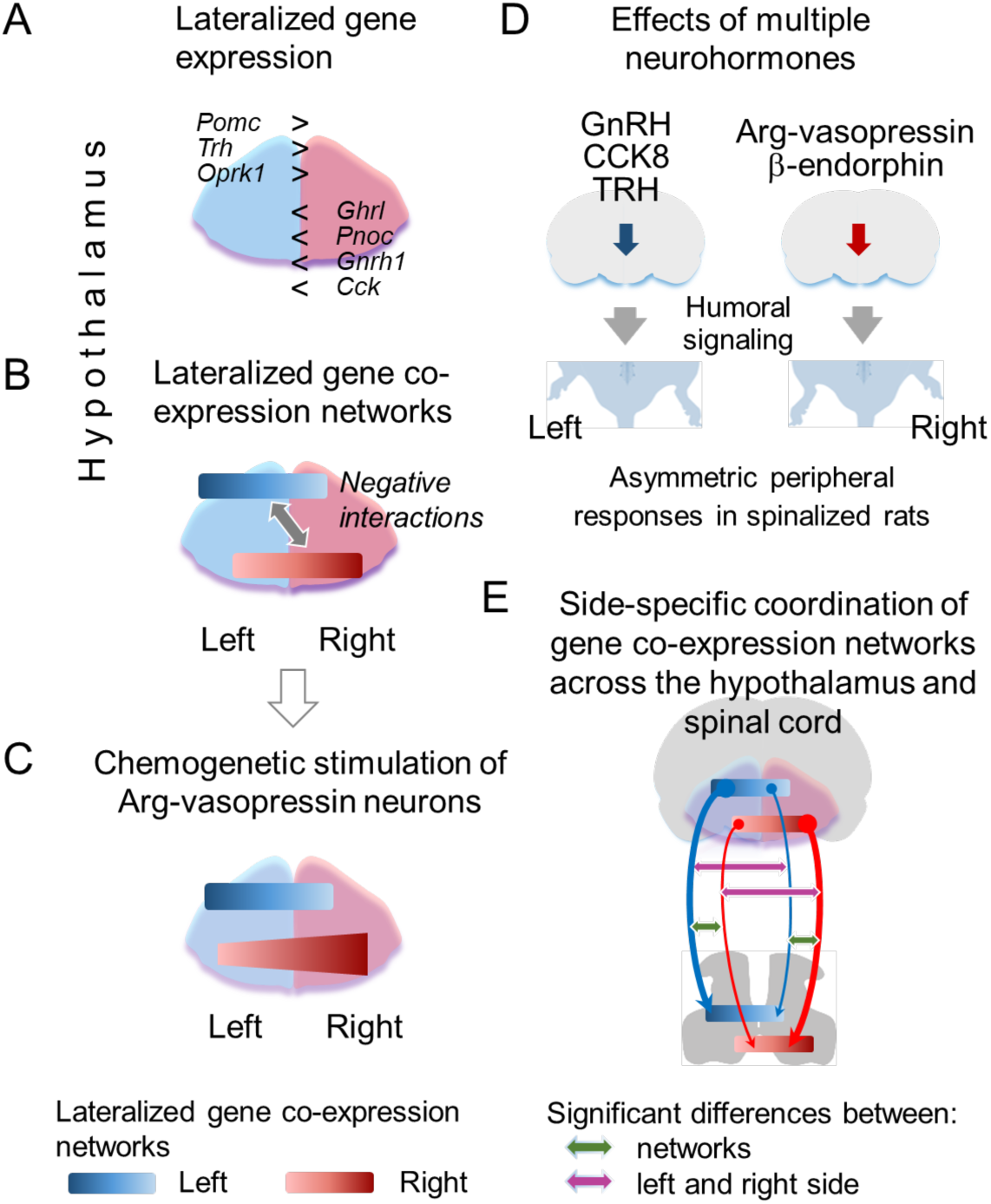
Schematic summary of findings and model of the lateralized neuroendocrine signaling from the hypothalamus to the spinal cord. (**A,B**) Lateralized expression of neurohormonal genes and lateralized co-expression networks of the neurohormonal and neuroplasticity-related genes in the hypothalamus. (**C**) The asymmetric effects of chemogenetic stimulation of AVP neurons on gene co-expression patterns: the right hypothalamic RdN is impacted. (**D**) Side-specific peripheral effects of multiple lateralized neurohormones: HL-PA was induced by intracisternal peptide administration in rats with complete spinal cord transection. The effects of Arg-vasopressin and β-endorphin were reported in the previous study (14). (**E**) Lateralized hypothalamic–spinal gene co-expression patterns. LdNs and RdNs exhibit distinct and reciprocally mirrored correlation profiles with spinal gene networks. These interactions are ipsilateral and differ between the left and right sides. Arrow shaft width roughly reflects the magnitude of left-right differences for both LdN and RdN. These patterns validate and extend previous results based on a limited set of neurohormonal genes (17). The findings depicted in (**C**-**E**) suggest a model for the regulation of left- and right side-specific peripheral processes by multiple lateralized neurohormones or their lateralized networks in the hypothalamus.

In addition to GnRH, we observed the asymmetric expression of ten other neurohormonal genes in the hypothalamus. Left-dominant genes included *Pomc*, *Trh*, and *Oprk1*, while right-dominant genes included *Cck*, *Pnoc*, *Gnrh1*, *Ghrl*, *Agtrap*, *Mas1*, and *Ren*. The left-sided dominance of *Pomc* aligns with the established role of β-endorphin—its peptide product—in mediating contralateral hindlimb postural effects following left-hemisphere injury (14). Similarly, asymmetric expression of *Cck* is consistent with evidence for lateralized behavioral effects of CCK-8 (78). The left-dominant expression of *Trh* mirrors reports of elevated TRH peptide levels in the left ventromedial, dorsomedial, and paraventricular nuclei of the human hypothalamus (79).

### Lateralized AVP coordination in the PVN

Another facet of hypothalamic lateralization lies in the microstructural asymmetry of AVP neuron coordination within the PVNpm suggested by stereology-based immunohistochemistry. This finding aligns with previous anatomical and neurochemical evidence indicating that the rostral (anteroventromedial) and caudal (posterodorsolateral) subdivisions of the PVNpm differ in cytoarchitecture (80) and peptidergic composition (47). Neurons in the caudal PVNpm are generally larger and more uniformly oriented, whereas the rostral PVNpm contains a higher proportion of oxytocin-expressing neurons (80–82). Moreover, ACTH- and β-endorphin–containing fibers and terminals are abundant in the rostral PVNpm but nearly absent in the caudal portion (83,84). The left-lateralized expression of *Pomc*, which encodes both ACTH and β-endorphin, suggests that asymmetric neurohormonal signaling—originating outside the PVNpm—may influence intra-nuclear AVP circuit organization. Specifically, neurohormonal input from the left hypothalamus may modulate AVP neuron interactions in a side-specific and subregion-specific manner.

The observed coordination between the right rostral and caudal PVNpm implies that AVP neurons in these subregions operate as an integrated functional network. In contrast, the decoupling of the left rostral PVNpm from this network suggests a distinct and possibly more autonomous role. Supporting this idea, patients with amyotrophic lateral sclerosis exhibit volume reductions in the left—but not right—anterior-superior hypothalamus, which includes the rostral PVN, implying a structurally and functionally less embedded role for this region in bilateral hypothalamic circuits (85).

A functional role of the asymmetry in AVP coordination remains to be revealed. However, it may be relevant for the lateralized effects of chemogenetic activation of AVP neurons, and side-specific encoding of hemispheric neural signals into neurohormonal outputs, particularly AVP itself.

### Lateralized gene co-expression networks and crosstalk between the hypothalamus and spinal cord

Gene co-expression analysis of all three rat cohorts revealed robust asymmetry in the hypothalamus, characterized by two distinct and lateralized networks: the LdN and the RdN. Within each network, genes were positively correlated, while interactions between the two were predominantly negative—suggesting antagonistic functions.

Chemogenetic stimulation of AVP neurons altered co-expression patterns of neurohormonal and neuroplasticity-related genes (**Figure 6**). Importantly, these effects were asymmetric, with the right hypothalamus showing the greatest changes. Such generalized effects may arise if neuroendocrine neurons are organized into integrated ensembles that respond collectively to perturbation of one element—and if these ensembles differ in architecture between the left and right hypothalamus.

Neuropeptides, neurohormones, and their receptors are typically expressed in distinct neuronal populations and across different hypothalamic subregions (18,19). These populations have been described to play defined roles mediated by a single cell type (86–88). This “labeled lines” mechanism applies to neurons, including the GnRH and TRH circuits that control pituitary hormone secretion. However, complex behaviors may be encoded by the combined activity of several classes of hypothalamic neurons. Combinatorial processing involving the coordinated activity of molecularly diverse neuroendocrine neurons has indeed been reported (86–88). In this model, “grouped ensemble” coding enables the coordination of multiple systems to execute integrated physiological and behavioral processes. Different neuroendocrine neurons communicate with each other (89–92) likely via receptors for multiple neuropeptides (88,93–95).

The observed gene co-expression patterns may arise from functional communication between populations of neuroendocrine neurons. Due to anatomical constraints that limit direct synaptic interactions among diverse spatially separated neuroendocrine neurons, neurohormone-mediated signaling is a more plausible mechanism by which the AVP circuit communicates with the broader neuroendocrine network than direct connectivity. A supporting evidence is the autocrine inhibition of phasic firing in vasopressin-releasing magnocellular neurons by endogenous dynorphins, which demonstrates how neuropeptides can shape network dynamics via local feedback (96).

Strikingly, the LdN and RdN also exhibited ipsilateral, asymmetric co-expression patterns between the hypothalamus and spinal cord, forming nearly mirror-symmetric correlation profiles: the LdN was more prominent on the left, and the RdN on the right (**Figure 6** in this study and (17)). These patterns were disrupted by UBI, with left UBI reversing the directionality of LdN-mediated asymmetry. In contrast, diagonal (contralateral) correlations between hypothalamic and spinal networks were weak and remained consistent across conditions. Notably, these co-expression patterns were observed in rats with fully transected thoracic spinal cords, indicating that the revealed hypothalamic–spinal coordination is not mediated by descending neural pathways but relies on humoral signaling mechanisms—consistent with the framework of the T-NES.

### Endocrine effects of lateralized neurohormones

Five lateralized hypothalamic neurohormones—GnRH, CCK-8, TRH, AVP, and β-endorphin—have been shown to induce side-specific peripheral effects via the humoral pathway when administered intracisternally, as demonstrated in this and previous studies (14). The side of the response depended on the specific neurohormone injected, indicating that these peptides can encode spatial information. After central administration, neurohormones may elicit asymmetric peripheral effects by acting on intrahypothalamic circuits or neural projections to the hypothalamus. These circuits may respond selectively to individual neurohormones or be activated by multiple peptides, but in a way that differentially enables either left- or right-sided output.

Previous studies demonstrated that AVP, β-endorphin, and Leu-enkephalin induced right hindlimb flexion, whereas Met-enkephalin, dynorphin, and κ-opioid receptor agonists caused left-sided flexion (6,14,15). Several of these peptides are processed from common precursors—Leu- and Met-enkephalins from proenkephalin, dynorphin and Leu-enkephalin from prodynorphin, and β-endorphin and Met-enkephalin from proopiomelanocortin. Therefore, side-specific outcomes may not solely depend on lateralized gene expression, but on asymmetric post-translational processing in the hypothalamus and also on lateralization of their receptors.

The findings of this study are novel and significantly expand our understanding of lateralized hypothalamic function. First, we demonstrate that both asymmetric expression of individual neurohormonal genes and lateralized gene co-expression networks are robust features of the hypothalamus under physiological conditions. These lateralized patterns were observed not only in naïve animals but also across two distinct experimental models—one physiological (chemogenetic AVP stimulation) and one pathological (unilateral brain injury). Previous molecular studies of hypothalamic lateralization were limited to highly perturbed states involving deep anesthesia, complete spinal cord transection, and focal brain lesions.

Second, the reproducibility and generalizability of these findings are supported by a large sample size—57 rats across three independent cohorts. To our knowledge, no prior studies of neuroendocrine lateralization have validated molecular asymmetries on this scale or across multiple conditions.

Third, our transcriptomic analysis encompassed nearly the full repertoire of hypothalamic neurohormonal genes, providing a comprehensive view of the lateralized neuroendocrine landscape. We report statistically significant lateralization for eight previously unreported genes, and confirm asymmetry in three additional genes across all three cohorts.

Fourth, we characterize distinct left- and right-dominant co-expression networks composed of neurohormonal genes. These networks were highly reproducible (with corrected permutation P values ranging from 0.05 to 9e-16), yet also flexible, showing cohort-specific gene composition and architecture. Importantly, these networks were inferred for the first time not only from an analysis of the correlational structure, but also from perturbation experiments. Chemogenetic stimulation of AVP neurons led to marked and asymmetric remodeling of these networks, predominantly in the right hypothalamus.

Finally, we show that three hypothalamic neurohormones—GnRH, CCK-8, and TRH—can each elicit side-specific peripheral effects when administered centrally. This provides the first direct evidence that these lateralized hormones can act through the humoral pathway to produce side-specific functional responses.

## Limitations

The first limitation concerns modeling lateralized neuroendocrine signaling under physiological settings. Ideally, the role of hypothalamic neuropeptides in encoding hemisphere-specific information into hormonal signals should be studied in minimally perturbed conditions. However, isolating neuroendocrine signaling from neural pathways necessitates complete spinal cord transection, as no alternative model currently allows for selective assessment of humoral signaling to the spinal cord.

Our HL-PA model provides a binary readout—left or right hindlimb flexion—serving as a proxy for lateralized deficits. While useful for acute experiments, it cannot be used in awake animals and does not capture the full spectrum of motor coordination and neurohormonal integration. To address the relevance of lateralized neuropeptide signaling in persistent physiological and pathophysiological contexts, future studies could employ subchronic experiments in spinalized, awake rats. These could integrate behavioral, electrophysiological, and biomechanical analyses—e.g., of body weight-supported stepping—to reveal effects of humoral signaling on motor output.

Mechanistic resolution of the endocrine pathway is the second limitation. While this study implicates hypothalamic neuropeptide signaling in side-specific regulation of peripheral function, the precise encoding and decoding mechanisms remain to be elucidated. These include identifying the central and peripheral targets of lateralized neuropeptides—potentially at the levels of hypothalamus, pituitary, spinal cord, or even spinal projections to hindlimb musculature. Further work is needed to resolve the identity and laterality of target motoneurons and interneurons, characterize relevant receptor distributions, and integrate molecular, electrophysiological, and functional data. A comprehensive mechanistic dissection would require a combination of pharmacological, genetic, and cell-type-specific manipulations. However, such resolution, while important, exceeds the realistic scope of this study.

The third limitation is that the portion of our results is based on transcriptional profiling, which may not fully reflect peptide abundance, post-translational processing, or biological activity. Lateralization is expected to manifest differently across regulatory layers—gene transcription, peptide processing, neuronal wiring, and circuit dynamics—all of which shape hypothalamic asymmetry. While spatially resolved peptidomics will be essential for future studies, the present analysis was designed to identify coherent transcriptional architectures and lateralized gene co-expression networks. Importantly, these networks cannot be captured by peptidomics and reflect an upstream regulatory layer.

Despite these limitations, our integrative approach—combining targeted neuronal stimulation, functional readouts, and transcriptomic network analysis—opens a previously unexplored dimension of neuroendocrine regulation and sets the stage for mechanistic investigations of lateralized hypothalamic control of peripheral function.

## CONCLUSIONS

This study demonstrates lateralized expression of neurohormonal genes and their co-expression networks in the hypothalamus. These networks are plastic, responding to circuit-level modulation in a side-specific manner, as shown by chemogenetic activation of AVP-producing neurons. We also suggest a novel form of lateralization—a microstructural asymmetry in the PVN—involving AVP, which acts as a side-specific neuroendocrine messenger between brain and periphery (14).

By analyzing three independent rat cohorts and employing a nearly comprehensive panel of hypothalamic neurohormonal genes regulating pituitary function, we obtained robust and reproducible evidence for the existence of a lateralized neuroendocrine architecture. Notably, multiple hypothalamic neurohormones—including TRH, GnRH, CCK-8, AVP, and opioid peptides (6,14–17)—were shown to elicit side-specific functional responses via humoral mechanisms supporting their role in endocrine regulation of left and right side-specific processes in the body.

These and other neurohormones may act either as individual lateralized regulators, or as components of integrated neurohormonal networks that coordinate left–right physiological processes. Such network-based integration could provide multi-layered control of lateralized outputs, independent of the specific action of any single neurohormone.

While the molecular and cellular mechanisms that encode lateralized signals remain to be fully elucidated, the present findings offer both molecular and functional evidence for a lateralized neuroendocrine axis linking the hypothalamus to peripheral endocrine targets. A deeper understanding of how the T-NES maintains left–right coordination—and how it may become dysregulated in disease—could ultimately lead to novel therapeutic strategies for conditions involving asymmetric impairments.

## Author contribution

H.W., O.N., A.L., E.L., R.F.F., S.M.S, N.L. and M.Z., performed injury, behavioral, morphological and molecular analysis. Y.U. and T.M. generated and guided experiments with transgenic rats. Y.K., D.S. and V.G. performed statistical analyses. H.W., N.L., I.L., A.G., M.H., J.S., M.Z. and G.B. planned the experiments, processed and discussed the data, and participated in manuscript preparation. M.Z. and G.B. conceived and supervised the project. G.B. wrote the manuscript. All authors worked with and commented on the manuscript.

## Acknowledgements

We are grateful to Ms. Karen Rich for assistance with the genotyping of AVP transgenic rats.

## GLOSSARY

### Asymmetry index

AI, the asymmetry index

AI_L/R_, the left / right asymmetry index computed as log_2_ (L / R), where L and R are values for the left and right sides. The AI_L/R_ was analyzed both for the gene expression levels and the resistance to stretch (W)

### Brain injury

UBI, a unilateral brain injury

### CNS areas

HPT, the hypothalamus

PVN, the paraventricular nucleus

PVNpm, the posterior magnocellular PVN

SpC, the spinal cord

### Correlation analysis

Coordination strength, magnitude of correlations (absolute value of the correlation coefficients) averaged across pairwise correlations

### Hindlimb postural asymmetry analysis

HL-PA, hindlimb postural asymmetry

MPA, the magnitude of postural asymmetry size (PAS) in mm

PAS, the postural asymmetry size in mm, with the direction of the asymmetry shown as negative values for left hindlimb flexion and positive values for right hindlimb flexion

### Musculo-articular resistance to stretching

W, the work in gm x mm for passive hindlimb musculo-articular resistance to stretching W_L-R_, the difference in the work applied to stretch the left (L) and right (R) hindlimbs

### Molecular analysis

LdN, the left dominant gene network consisting of genes with the asymmetry index AI_L/R_ > 0

RdN, the right dominant gene network consisting of genes with the asymmetry index AI _L/R_ < 0

LM, the left module constituted by gene transcripts of the left half of the hypothalamus or the spinal cord

RM, the right module constituted by gene transcripts of the right half of the hypothalamus or the spinal cord

### Neurohormones

AVP, Arg-vasopressin

CCK-8, cholecystokinin-8

GnRH, gonadotropin-releasing hormone

TRH, thyrotropin-releasing hormone

### Statistical terms

95% HPD, the highest posterior density credible interval, within which an unobserved parameter value falls with 95% probability. This is a Bayesian analog of 95% confidence interval

Significant asymmetry, 95% HPD does not include zero and adjusted P-value < 0.05 Contrast, the median of one group minus the median of another group

## Declaration of generative AI and AI-assisted technologies in the writing process

During the preparation of this work the author (G.B.) used ChatGPT4o in order to polish text. After using this service, the author reviewed and edited the content as needed and takes full responsibility for the content of the publication.

## VISUAL SUMMARY

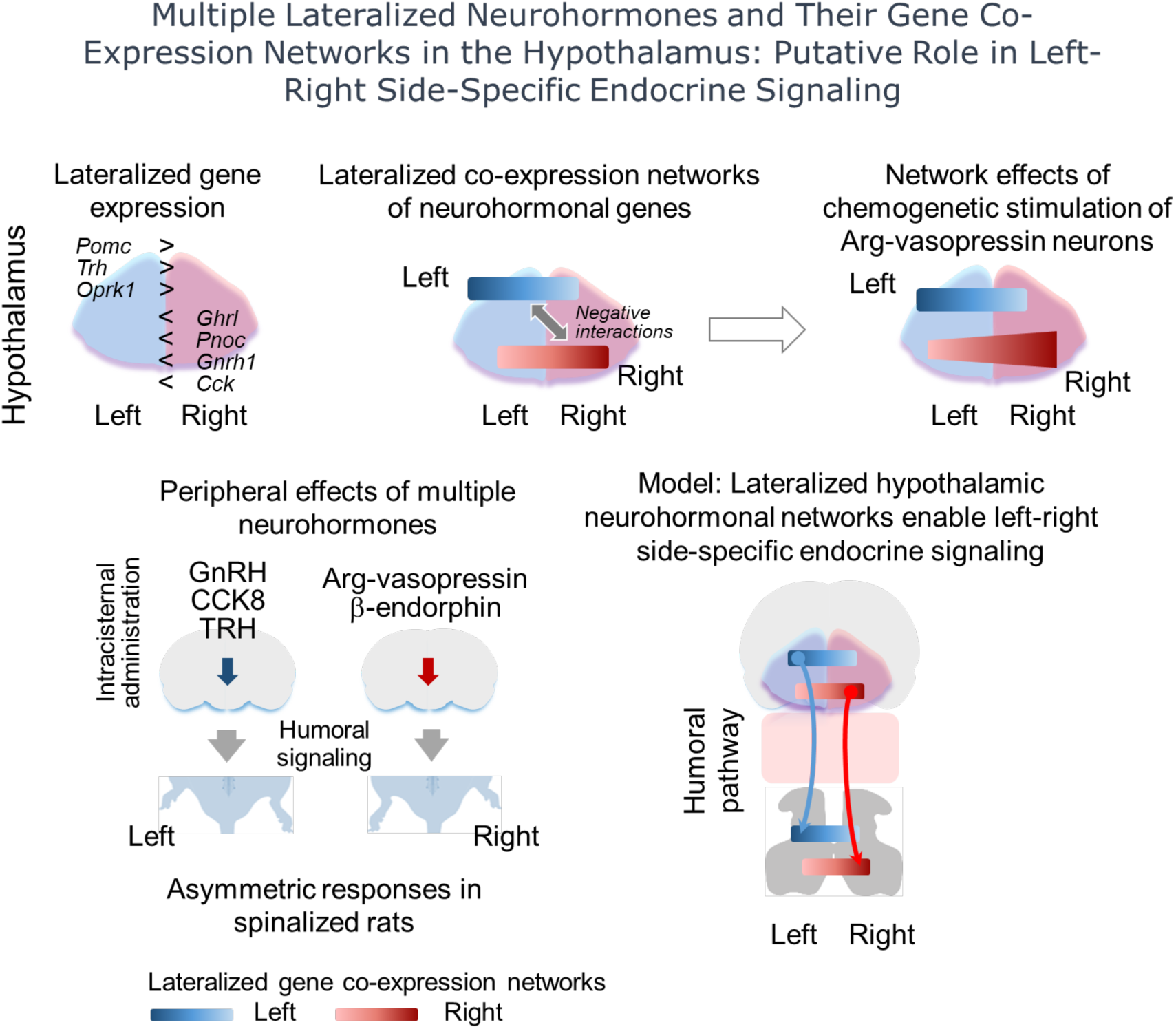

## SUPPLEMENTARY FIGURES AND TABLES

**Supplementary Figure 1.**
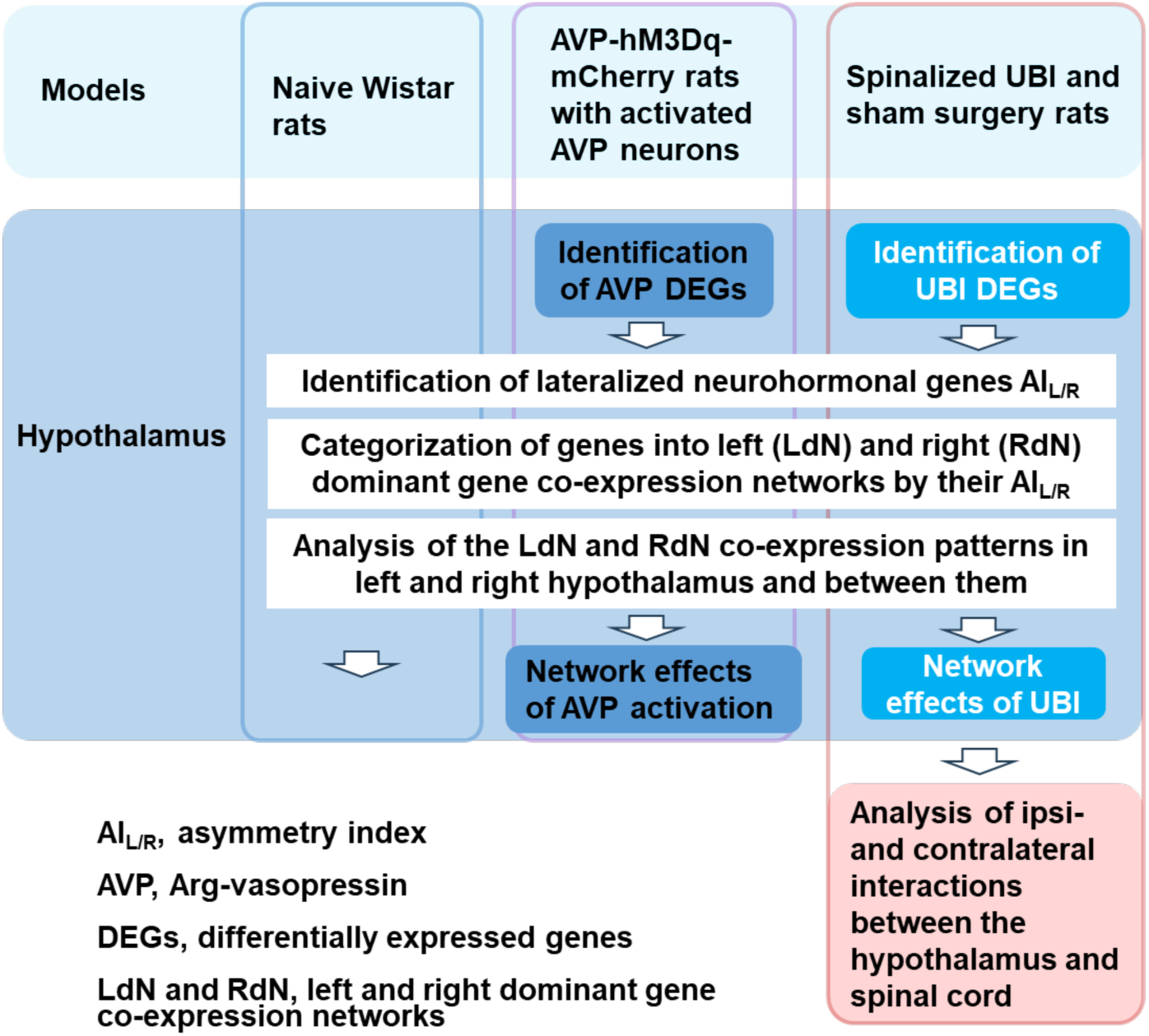
Design of gene expression and co-expression analysis. Rat cohorts analyzed included i) naïve Wistar rats (n = 11); transgenic AVP-hM3Dq-mCherry rats expressing the human M1 receptor gene under vasopressin promoter in the hypothalamus which were treated with CNO (n = 11 rats) to activate the transgene, or with vehicle control (n = 11); and the spinalized UBI (n = 12) and sham surgery (n = 11) rats. Expression of the neurohormonal genes, neuropeptides and their receptors genes, and neuroplasticity-related genes was analyzed in the left and right hypothalamus (n = 47 - 48 genes; **Supplementary Table 1**), and expression of the neuropeptides and their receptors genes and neuroplasticity-related genes in the left and right lumbar spinal cord (n = 27 genes; **Supplementary Table 2**). In the hypothalamus, differentially expressed genes (DEGS) in the transgenic rats treated with CNO, and in UBI rats, and genes with lateralized expression in three cohorts were searched for. Genes were categorized to the lateralized gene co-expression networks based on their asymmetry index AI_L/R_ = log_2_[L/R], where L and R are the median expression levels in the left and right hypothalamus or the left and right spinal cord. Genes with higher expression either on the left or right side were defined as genes of the left dominant network (LdN; AI_L/R_ > 0) or right dominant network (RdN; AI_L/R_ < 0), respectively. The patterns of interactions between LdN and RdN were analyzed as the coordination strength (magnitude of the correlations or the absolute value of the correlation coefficient averaged over pairwise correlations) and assessed as the proportion of positive correlations. The patterns were compared between the networks on the left and right side, between the sides, and across analyzed areas and tissues.

**Supplementary Figure 2.**
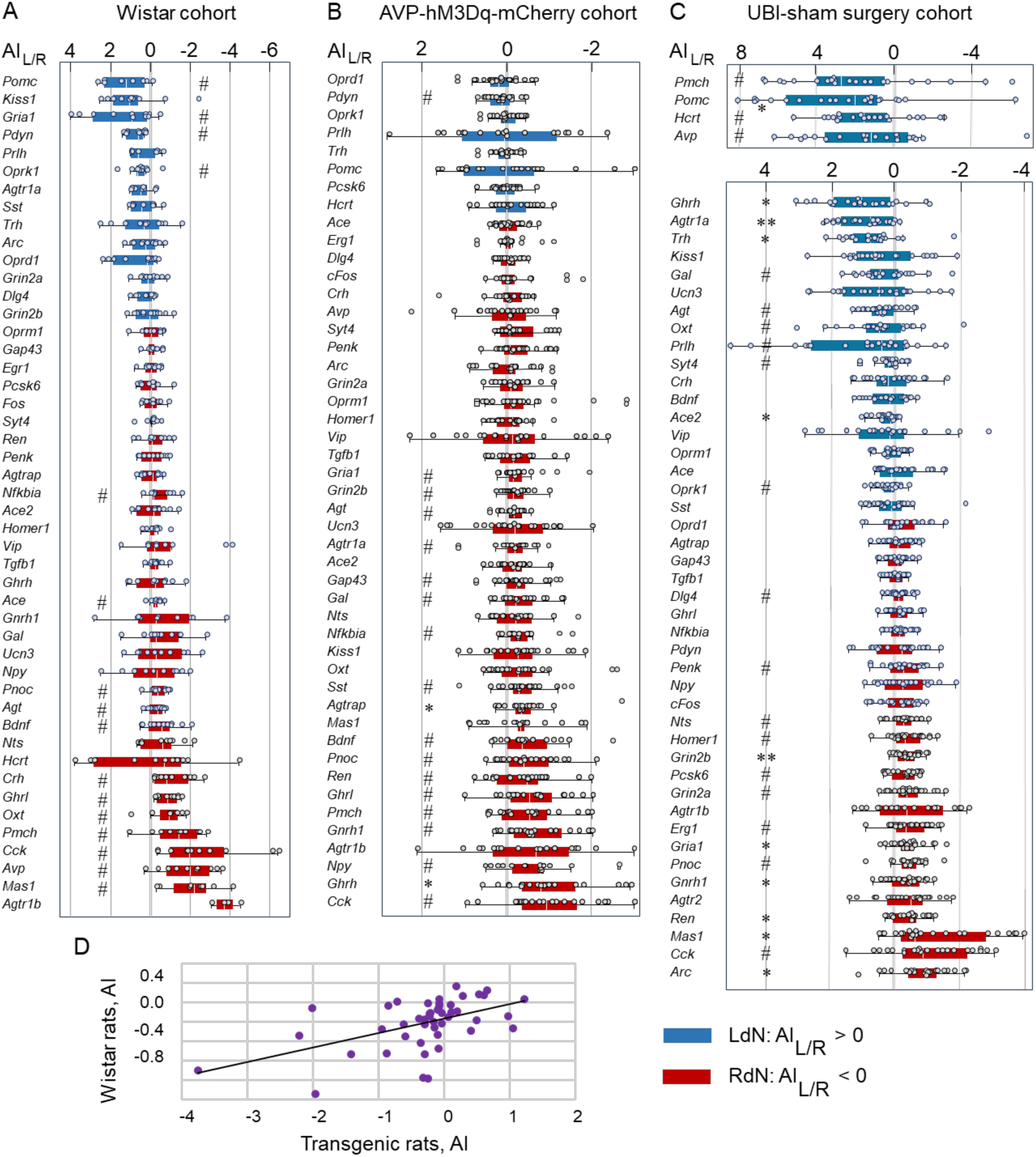
(A-C) Lateralized expression of the neurohormonal and neuroplasticity-related genes in the rat hypothalamus and gene categorization into the left and right co-expression networks LdN and RdN, respectively. The AI_L/R_ is shown for each gene analyzed. (**A**) Naïve Wistar rats (n = 11). (**B**) Transgenic AVP-hM3Dq-mCherry rats; there were no differences in the AI_L/R_ between female and male subgroups, and between CNO (n = 11) and vehicle (n = 11) treated rats in the AI_L/R_; therefore, they were combined (n = 22) for statistical analysis of the asymmetry of expression of individual genes (Mann–Whitney test followed by a Bonferroni correction). (**C**) The UBI and sham surgery cohorts. There were no differences in the AI_L/R_ between UBI (n = 12) and sham surgery (n = 11) groups; therefore, they were combined (n = 23) for statistical analysis of the asymmetry of expression of individual genes (Mann–Whitney test followed by a Bonferroni correction). Data are presented as boxplots with median and hinges representing the first and third quartiles, and whiskers extending from the hinge to the highest/lowest value that lies within the 1.5 interquartile range of the hinge. Data for individual rats are shown as circles. Blue boxes denote genes with AI_L/R_ > 0, and pink boxes with AI_L/R_ < 0 that were defined as the LdN and RdN genes, respectively. A two-sided Wilcoxon signed rank exact test was used to assess significance of lateralization: *, P < 0.05 and **, P < 0.01 after Bonferroni correction (n = 47 genes for Wistar cohort and AVP-hM3Dq-mCherry transgenic cohort; n = 48 for the UBI cohort); #, genes showing nominally significant AI_L/R_ (P < 0.05; not adjusted). Data for expression of 28 genes in the UBI-sham surgery cohort were taken from our previous study.(17) (**D**) Pearson correlation in the AI_L/R_ between the Wistar cohort and the transgenic AVP-hM3Dq-mCherry cohort. The AI_L/R_ values are shown for the median expression levels of 47 genes. Pearson correlation coefficient: 0.52, P = 2e-04; Spearman’s rank correlation coefficient: 0.53, P = 1.2e-04.

**Supplementary Figure 3.**
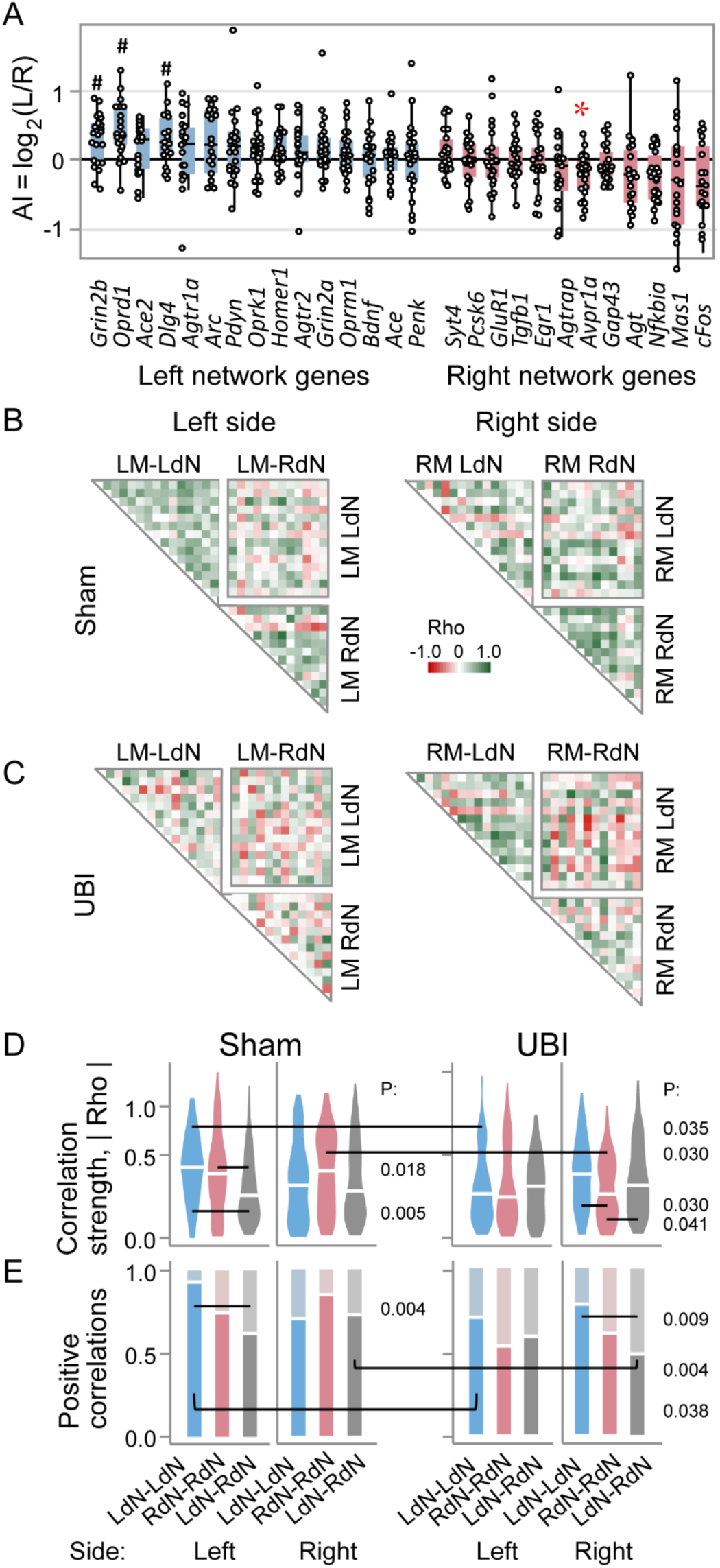
Gene expression patterns in the lumbar spinal cord. Analysis of the LdN and RdN gene networks and the UBI effects on the LdN and RdN. Gene expression was analyzed in the spinal cord LM and RM of the left sham surgery (n = 11) and left UBI (n = 12) spinalized rats. Spinal cords were transected prior to brain injury. (**A**) Gene categorization using the AI_L/R_ into the LdN (AI_L/R_ > 0) and RdN (AI_L/R_ < 0). There were no differences in the AI_L/R_ between sham surgery and UBI groups; therefore, they were combined (n = 22) for statistical analysis. The median AI_L/R_ is shown for the combined left sham surgery and the left UBI group. Data are presented as boxplots with median and hinges representing the first and third quartiles, and whiskers extending from the hinge to the highest/lowest value that lies within the 1.5 interquartile range of the hinge. Wilcoxon signed-rank test followed by Bonferroni multiple testing correction: *, P adj < 0.05; #, P ≤ 0.05 (not adjusted). (**B,C**) Heatmaps for Spearman’s rank coefficients for pairwise gene-gene correlations in the LM and RM of sham surgery and UBI groups. (**D,E**) Patterns of intra-modular correlations internal for each the LdN (LdN-LdN) and RdN (RdN-RdN), and between the networks (LdN-RdN). Strengths of pairwise correlations (correlation coefficient |Rho|) are presented as violin plots with white line indicating the coordination strength (the absolute Rho value averaged across correlations) and the proportion of positive correlations for the LM and RM are shown for the sham surgery and UBI groups. The correlation patterns were compared within each LM and RM and each pattern was compared between the modules, and between the sham surgery and UBI groups. Shown are P values that were determined by permutation testing with Benjamini-Hochberg family-wise multiple test correction.

**Supplementary Figure 4.**
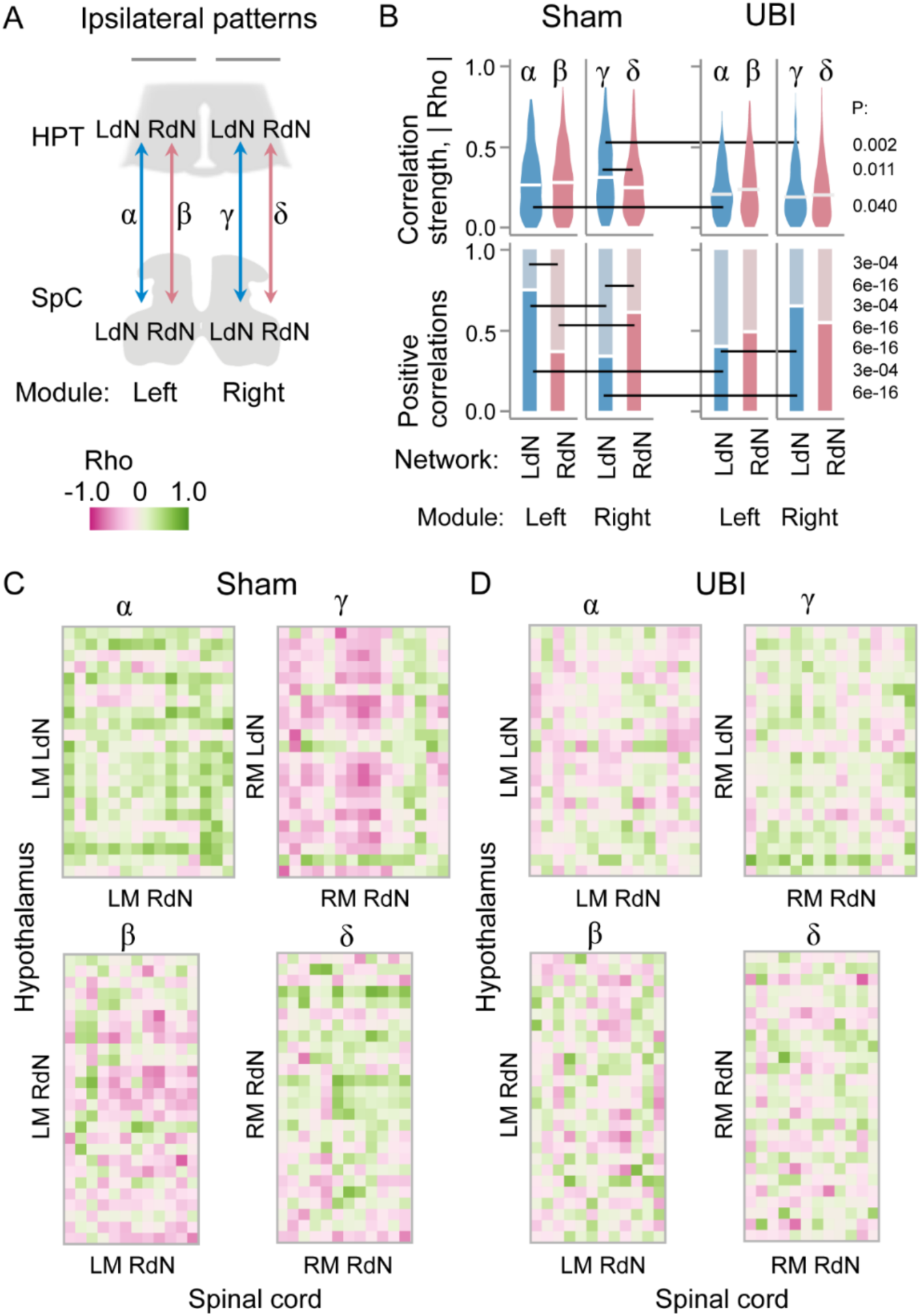
Ipsilateral correlation patterns of the LdN and RdN between the hypothalamus and lumbar spinal cord. The effects of UBI. (**A**) Analyzed patterns of the ipsilateral pairwise gene-gene Spearman rank correlations between the hypothalamus (HPT) and spinal cord (SpC) on the left side (α and β) and on the right side (γ and δ). (**B**) The coordination strength and the proportion of positive correlations for the correlation patterns depicted in **A**. The correlation patterns were compared between the LdN and RdN (α *vs*. β; γ *vs*. δ); each of them between the left and right modules (α *vs*. γ; β *vs*. δ), and all four patterns individually between UBI and sham surgery groups. Shown are P values that were determined by permutation testing with Benjamini-Hochberg family-wise multiple test correction. (**C**,**D**) Heatmaps for Spearman’s rank coefficients for pairwise gene-gene correlations for the LM and RM in both the sham surgery and UBI groups.

**Supplementary Figure 5.**
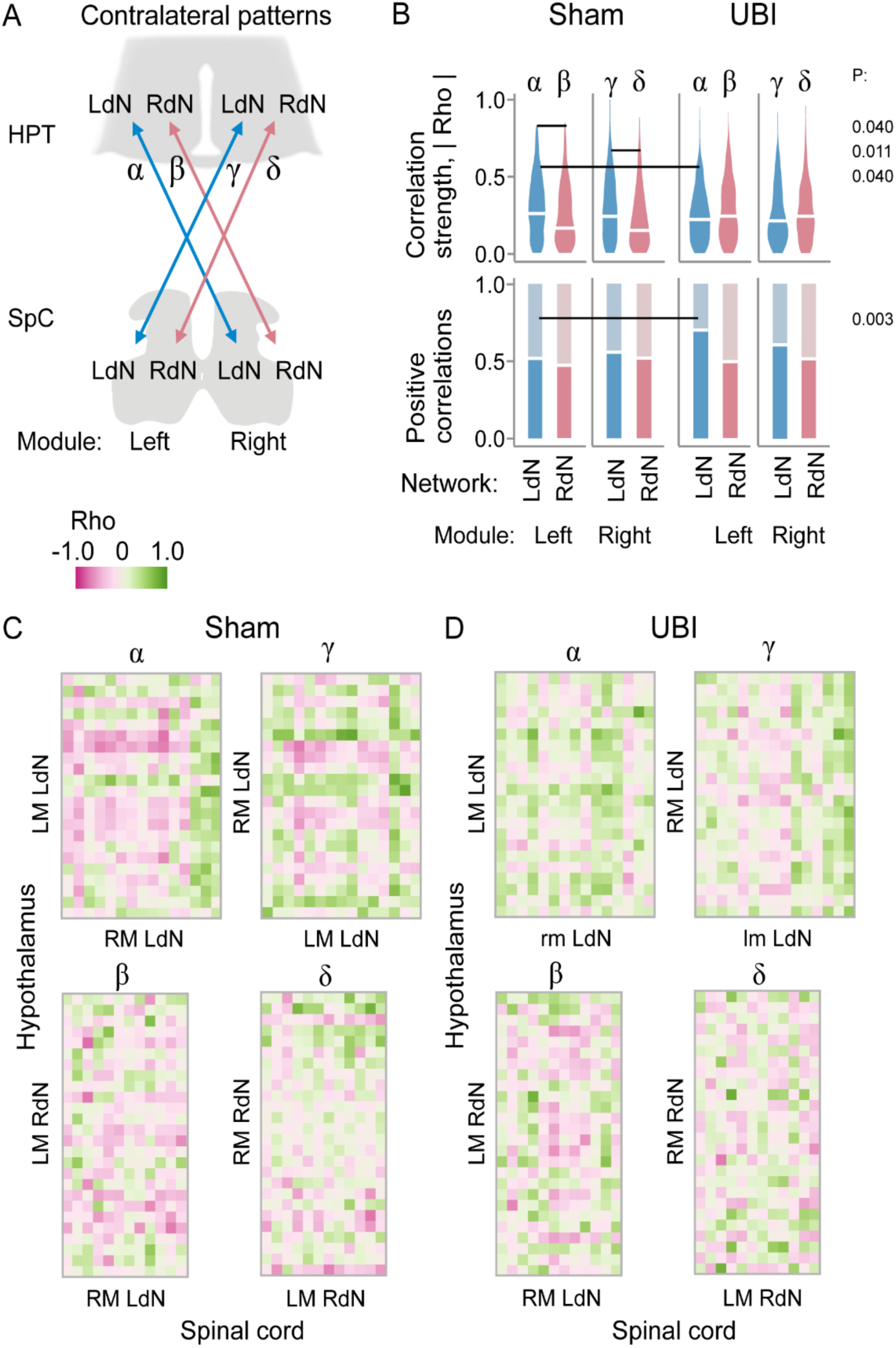
Contralateral correlation patterns of the LdN and RdN between the hypothalamus and lumbar spinal cord. The effects of UBI. (**A**) Analyzed patterns of the contralateral pairwise gene-gene Spearman rank correlations between the left hypothalamus and right spinal cord (α and β), and between right hypothalamus and left spinal cord (γ and δ). (**B**) The coordination strength and the proportion of positive correlations for the correlation patterns depicted in **A**. The correlation patterns were compared between the LdN and RdN (α *vs*. β; γ *vs*. δ); each of them between the left and right modules (α *vs*. γ; β *vs*. δ), and all four patterns individually between UBI and sham surgery groups. Shown are P values that were determined by permutation testing with Benjamini-Hochberg family-wise multiple test correction. (**C**,**D**) Heatmaps for Spearman’s rank coefficients for pairwise gene-gene correlations for the LM and RM in both the sham surgery and UBI groups.

**Supplementary Figure 6.**
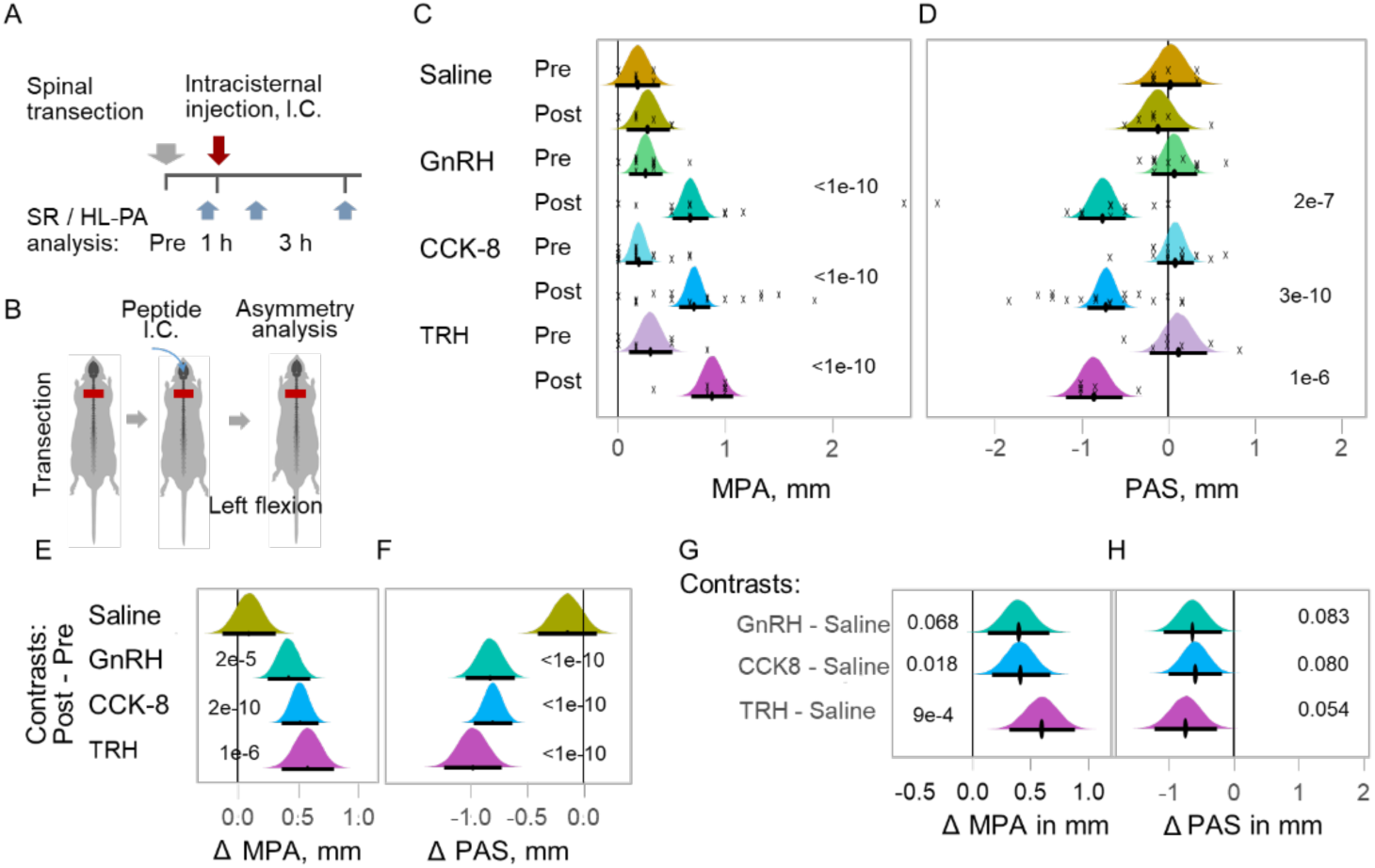
HL-PA induced by neurohormones administered into the cisterna magna of rats with completely transected thoracic spinal cords. (**A,B**) Experimental design. The spinal cord was transected at the T2-T4 level that was followed by intracisternal administration of peptides or saline. HL-PA was manifested as flexion of the left or right hindlimb. Data acquired before (Pre) and 3 h after (Post) injection are shown. Significance of the peptide effects was essentially the same for the 1 h-time point. (**C**) The magnitude of postural asymmetry (MPA; or absolute values of HL-PA) measured in millimeters (mm). (**D**) The size of postural asymmetry (PAS) measured in millimeters (mm). Negative and positive PAS values are assigned to rats with left and right hindlimb flexion, respectively. The MPA and PAS values for individual rats are indicated by crosses. (**E,F**) The contrasts between Post and Pre values for both MPA and PAS. (**G,H**) The contrasts in MPA and PAS between the peptide and saline groups denoted as ΔMPA and ΔPAS, respectively. The median (represented as black circles), 95% HPD (black lines), and posterior density (colored distribution) from Bayesian regression are used to plot the MPA and PAS and contrasts. Asymmetry and contrasts among the groups were deemed significant, with a 95% HPD not encompassing zero and adjusted P values of ≤ 0.05. Adjusted P values are presented numerically on the plots for significant asymmetries (in **C** and **D**), and for significant contrasts (in **E**-**H**). Rats received saline (n = 6); GnRH at doses 0.1 (n = 3 rats), 1 (n = 6 rats), and 50 (n = 2 rats) ng/rat; CCK-8 at doses 0.1 (n = 2 rats), 1 (n = 6 rats), 10 (n = 2 rats), and 30 (n = 7 rats) ng/rat; or TRH at dose 1 ng/rat (n = 7 rats). No significant differences between the doses for GnRH were revealed and the data were combined (dose was included as covariate in statistical analysis; see Materials and Methods). The same holds true for CCK-8.

**Supplementary Table 1.** Genes coding for hypothalamic releasing and inhibitory hormones, neuropeptides and neuroplasticity related genes and PCR probes for analysis of their expression levels (Bio-Rad Laboratories, CA, USA).

| Gene symbol | Gene name | Assay ID | Channel |
| --- | --- | --- | --- |
| *Ace* | *Angiotensin I converting enzyme* | Rn00561094_m1 | FAM |
| *Ace2* | *Angiotensin converting enzyme 2* | qRnoCIP0029204 | FAM |
| *Actb* | *Actin beta* | qRnoCIP0050804 | FAM |
| *Agt* | *Angiotensinogen* | qRnoCEP0045486 | TEX 615 |
| *Agtr1a* | *Angiotensin II receptor type 1a* | qRnoCIP0046446 | HEX |
| *Agtr1b* | *Angiotensin II receptor type 1b* | qRnoCEP0027560 | FAM |
| *Agtr2* | *Angiotensin II receptor type 2* | qRnoCEP0029382 | FAM |
| *Agtrap* | *Angiotensin II receptor associated protein* | qRnoCEP0026882 | TEX 615 |
| *Avp* | *Arginine Vasopressin* | qRnoCEP0023611 | FAM |
| *Arc* | *Activity-regulated cytoskeleton associated protein* | qRnoCEP0027389 | FAM |
| *Bdnf* | *Brain derived neurotrophic factor* | qRnoCEP0026843 | FAM |
| *Cck* | *Cholecystokinin* | qRnoCEP0030064 | FAM |
| *cFos* | *Fos proto-oncogene* | qRnoCEP0024078 | FAM |
| *Crh* | *Corticotropin releasing hormone* | qRnoCEP0026204 | FAM |
| *Dlg4* | *Discs large MAGUK scaffold protein 4* | qRnoCIP0026242 | FAM |
| *Egr1* | *Early growth response 1* | qRnoCEP0022872 | FAM |
| *Gal* | *Galanin and GMAP prepropeptide* | qRnoCIP0029109 | FAM |
| *Gap43* | *Growth associated protein 43* | qRnoCIP0027599 | FAM |
| *Gapdh* | *Glyceraldehyde-3-phosphate dehydrogenase* | qRnoCIP0050838 | FAM |
| *Ghrh* | *Growth hormone releasing hormone* | qRnoCIP0029554 | FAM |
| *Ghrl* | *Ghrelin and obestatin prepropeptide* | qRnoCEP0024487 | FAM |
| *Gnrh1* | *Gonadotropin releasing hormone 1* | qRnoCIP0025960 | FAM |
| *Gria1* | *Glutamate Ionotropic Receptor AMPA Type Subunit 1* | qRnoCIP0030725 | FAM |
| *Grin2a* | *Glutamate ionotropic receptor NMDA type subunit 2a* | qRnoCIP0025244 | FAM |
| *Grin2b* | *Glutamate ionotropic receptor NMDA type subunit 2b* | qRnoCIP0023973 | FAM |
| *Hcrt* | *Hypocretin neuropeptide precursor* | qRnoCEP0025559 | FAM |
| *Homer1* | *Homer scaffold protein 1* | qRnoCEP0023985 | FAM |
| *Kiss1* | *Kiss-1 metastasis suppressor* | qRnoCEP0045127 | FAM |
| *Mas1* | *Mas1 proto-oncogene G-protein-coupled receptor* | qRnoCEP0031088 | FAM |
| *Nfkbia* | *NFkB inhibitor alpha* | qRnoCEP0026759 | FAM |
| *Npy* | *Neuropeptide Y* | qRnoCEP0031211 | FAM |
| *Nts* | *Neurotensin* | qRnoCIP0026101 | FAM |
| *Oprd1* | *Opioid receptor delta 1* | qRnoCEP0029668 | FAM |
| *Oprk1* | *Opioid receptor kappa 1* | qRnoCIP0029310 | FAM |
| *Oprm1* | *Opioid receptor mu 1* | qRnoCEP0024902 | FAM |
| *Oxt* | *Oxytocin* | qRnoCEP0031434 | HEX |
| *Pcsk6* | *Proprotein convertase subtilisin/kexin type 6* | qRnoCIP0045340 | FAM |
| *Pdyn* | *Prodynorphin* | qRnoCEP0025357 | FAM |
| *Penk* | *Proenkephalin* | qRnoCEP0029455 | FAM |
| *Pmch* | *Pro-melanin concentrating hormone* | qRnoCEP0047681 | FAM |
| *Pnoc* | *Prepronociceptin* | qRnoCEP0029972 | FAM |
| *Pomc* | *Proopiomelanocortin* | pRnoCIP0024350 | FAM |
| *Prlh* | *Prolactin releasing hormone* | qRnoCEP0026951 | FAM |
| *Ren* | *Renin* | qRnoCIP0030552 | FAM |
| *Sst* | *Somatostatin* | qRnoCEP0031118 | FAM |
| *Syt4* | *Synaptotagmin 4* | qRnoCIP0029728 | FAM |
| *Tgfb1* | *Transforming growth factor beta 1* | qRnoCIP0031022 | FAM |
| *Trh* | *Thyrotropin releasing hormone* | qRnoCEP0026268 | FAM |
| *Ucn3* | *Urocortin 3* | qRnoCEP0024166 | FAM |
| *Vip* | *Vasoactive intestinal peptide* | qRnoCIP0026148 | FAM |

**Supplementary Table 2.** Genes coding for spinal cord genes, and PCR probes for analysis of their expression levels (Bio-Rad Laboratories, CA, USA).

| Gene symbol | Gene name | Assay ID | Channel |
| --- | --- | --- | --- |
| *Ace* | *Angiotensin I converting enzyme* | Rn00561094_m1 | FAM |
| *Ace2* | *Angiotensin converting enzyme 2* | qRnoCIP0029204 | FAM |
| *Actb* | *Actin beta* | qRnoCIP0050804 | FAM |
| *Agt* | *Angiotensinogen* | qRnoCEP0045486 | TEX 615 |
| *Agtr1a* | *Angiotensin II receptor type 1a* | qRnoCIP0046446 | HEX |
| *Agtr2* | *Angiotensin II receptor type 2* | qRnoCEP0029382 | FAM |
| *Agtrap* | *Angiotensin II receptor associated protein* | qRnoCEP0026882 | TEX 615 |
| *Arc* | *Activity-regulated cytoskeleton associated protein* | qRnoCEP0027389 | FAM |
| *Avpr1a* | *Arginine vasopressin receptor 1a* | qRnoCEP0023750 | FAM |
| *Bdnf* | *Brain derived neurotrophic factor* | qRnoCEP0026843 | FAM |
| *cFos* | *Fos proto-oncogene* | qRnoCEP0024078 | FAM |
| *Dlg4* | *Discs large MAGUK scaffold protein 4* | qRnoCIP0026242 | FAM |
| *Egr1* | *Early growth response 1* | qRnoCEP0022872 | FAM |
| *Gap43* | *Growth associated protein 43* | qRnoCIP0027599 | FAM |
| *Gapdh* | *Glyceraldehyde-3-phosphate dehydrogenase* | qRnoCIP0050838 | FAM |
| *Gria1* | *Glutamate Ionotropic Receptor AMPA Type Subunit 1* | qRnoCIP0030725 | FAM |
| *Grin2a* | *Glutamate ionotropic receptor NMDA type subunit 2a* | qRnoCIP0025244 | FAM |
| *Grin2b* | *Glutamate ionotropic receptor NMDA type subunit 2b* | qRnoCIP0023973 | FAM |
| *Homer1* | *Homer scaffold protein 1* | qRnoCEP0023985 | FAM |
| *Mas1* | *Mas1 proto-oncogene G-protein-coupled receptor* | qRnoCEP0031088 | FAM |
| *Nfkbia* | *NFkB inhibitor alpha* | qRnoCEP0026759 | FAM |
| *Oprd1* | *Opioid receptor delta 1* | qRnoCEP0029668 | FAM |
| *Oprk1* | *Opioid receptor kappa 1* | qRnoCIP0029310 | FAM |
| *Oprm1* | *Opioid receptor mu 1* | qRnoCEP0024902 | FAM |
| *Pcsk6* | *Proprotein convertase subtilisin/kexin type 6* | qRnoCIP0045340 | FAM |
| *Pdyn* | *Prodynorphin* | qRnoCEP0025357 | FAM |
| *Penk* | *Proenkephalin* | qRnoCEP0029455 | FAM |
| *Syt4* | *Synaptotagmin 4* | qRnoCIP0029728 | FAM |
| *Tgfb1* | *Transforming growth factor beta 1* | qRnoCIP0031022 | FAM |

**Supplementary Table 3.** Total number of AVP-stained cells in the posterior magnocellular subdivision of the hypothalamic paraventricular nucleus (PVNpm). Data represent mean (SD). Separate estimates were made in the rostral and caudal parts of this subdivision. As there were no effects of treatment on cell counts, data from animals in four groups (control, sham-UBI, left-UBI and right-UBI) were pooled together (n =21). No significant effects of lateralization on the number of AVP cells in the PVNpm and its rostral and caudal parts were detected (two-way ANOVA and paired *t*-test).

| PVN subdivision | Left side | Right side | P |
| --- | --- | --- | --- |
| Total PVNpm | 977 (304) | 968 (321) | n.s. |
| Rostral PVNpm | 414 (167) | 413 (142) | n.s. |
| Caudal PVNpm | 562 (217) | 556 (213) | n.s. |

**Supplementary Table 4.** The number of rats used in analysis of the effects of neuropeptides on HL-PA and hindlimb resistance to stretch.

| Treatment | Dose<br>ng / rat | The number<br>of rats |
| --- | --- | --- |
| Saline | - | 6 |
| TRH | 1.0 | 7 |
| GnRH | 0.1 | 3 |
| GnRH | 1.0 | 6 |
| GnRH | 50 | 2 |
| CCK-8 | 0.1 | 2 |
| CCK-8 | 1.0 | 6 |
| CCK-8 | 10 | 2 |
| CCK-8 | 30 | 7 |

### Key resources table

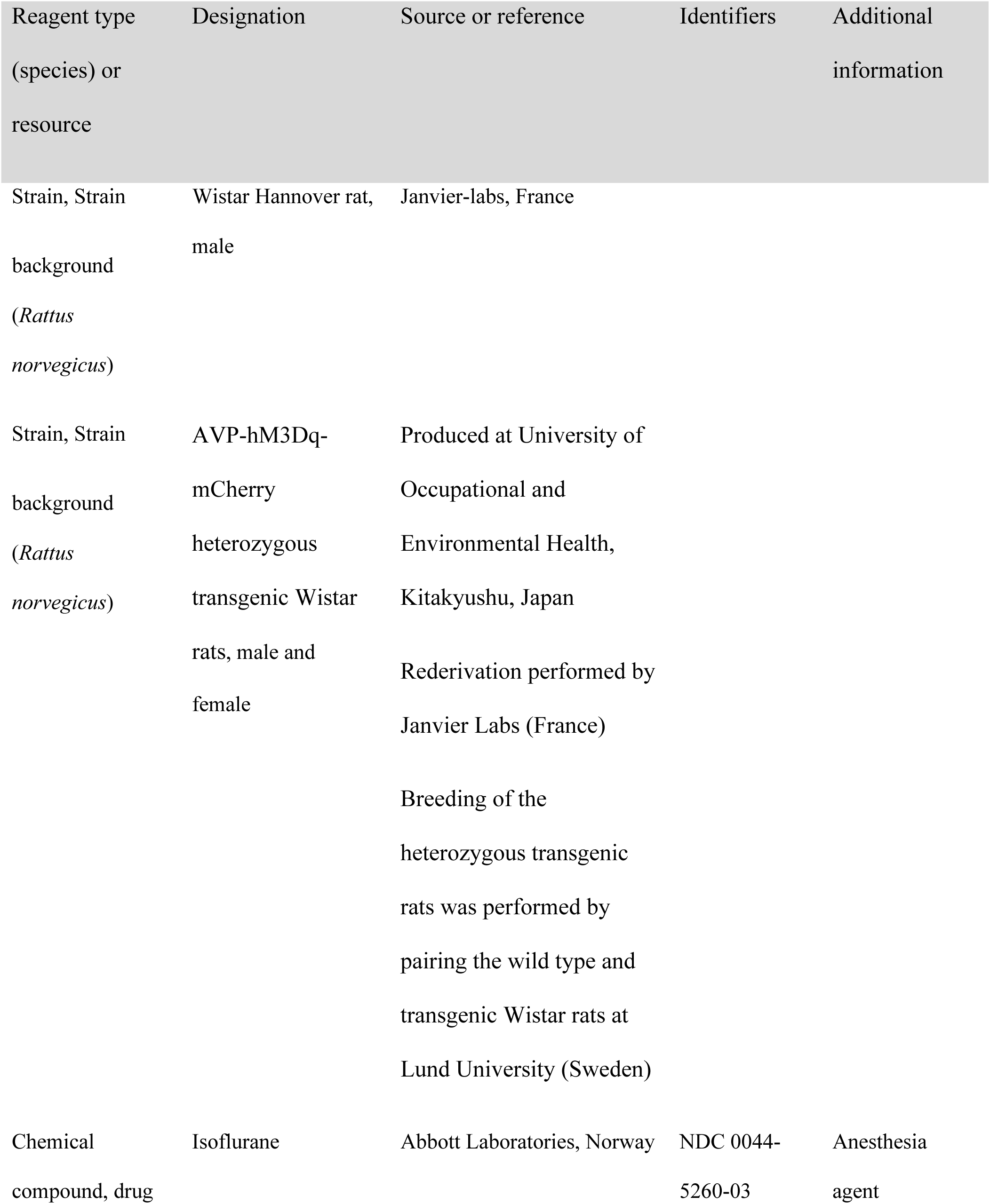
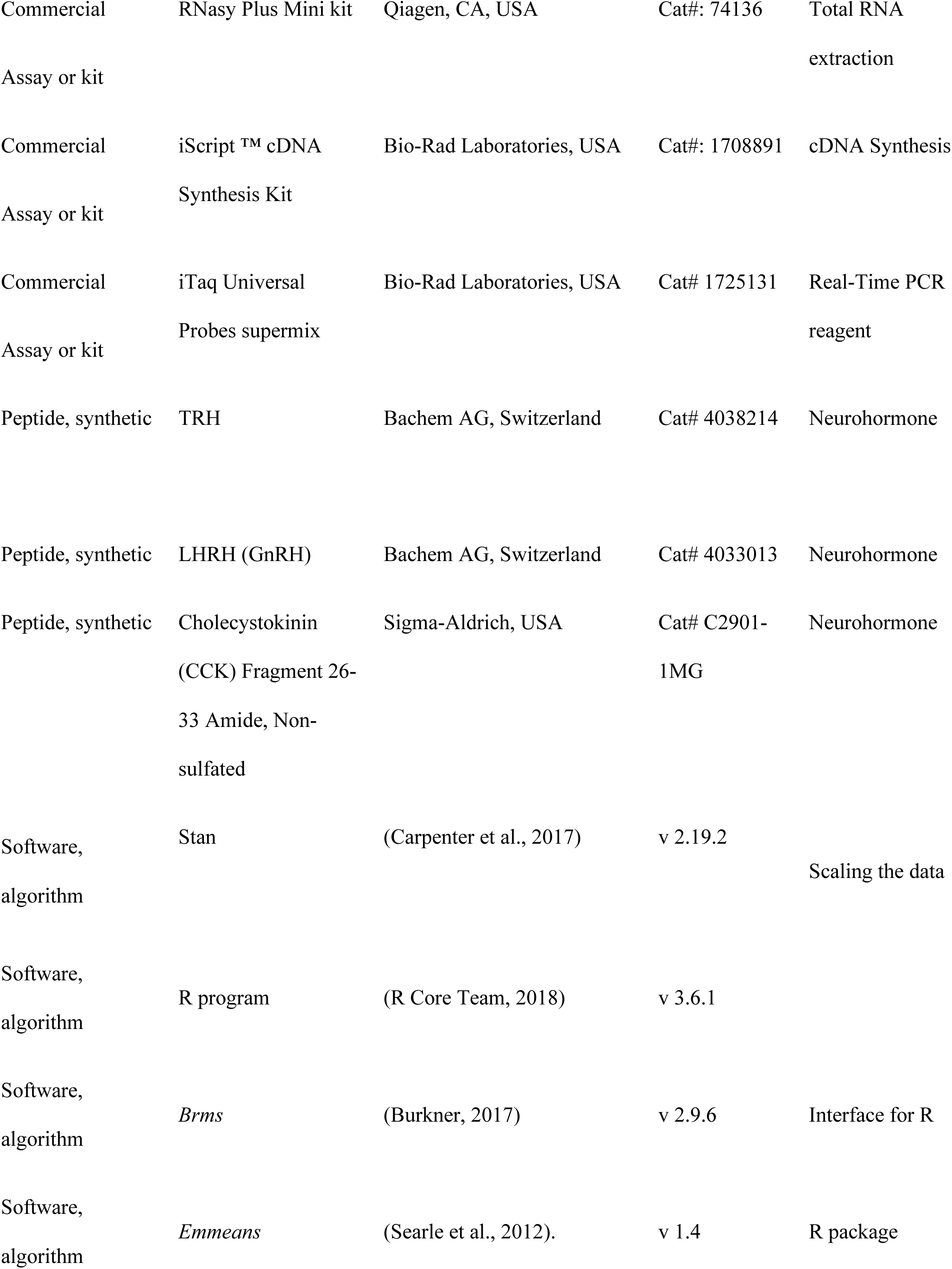
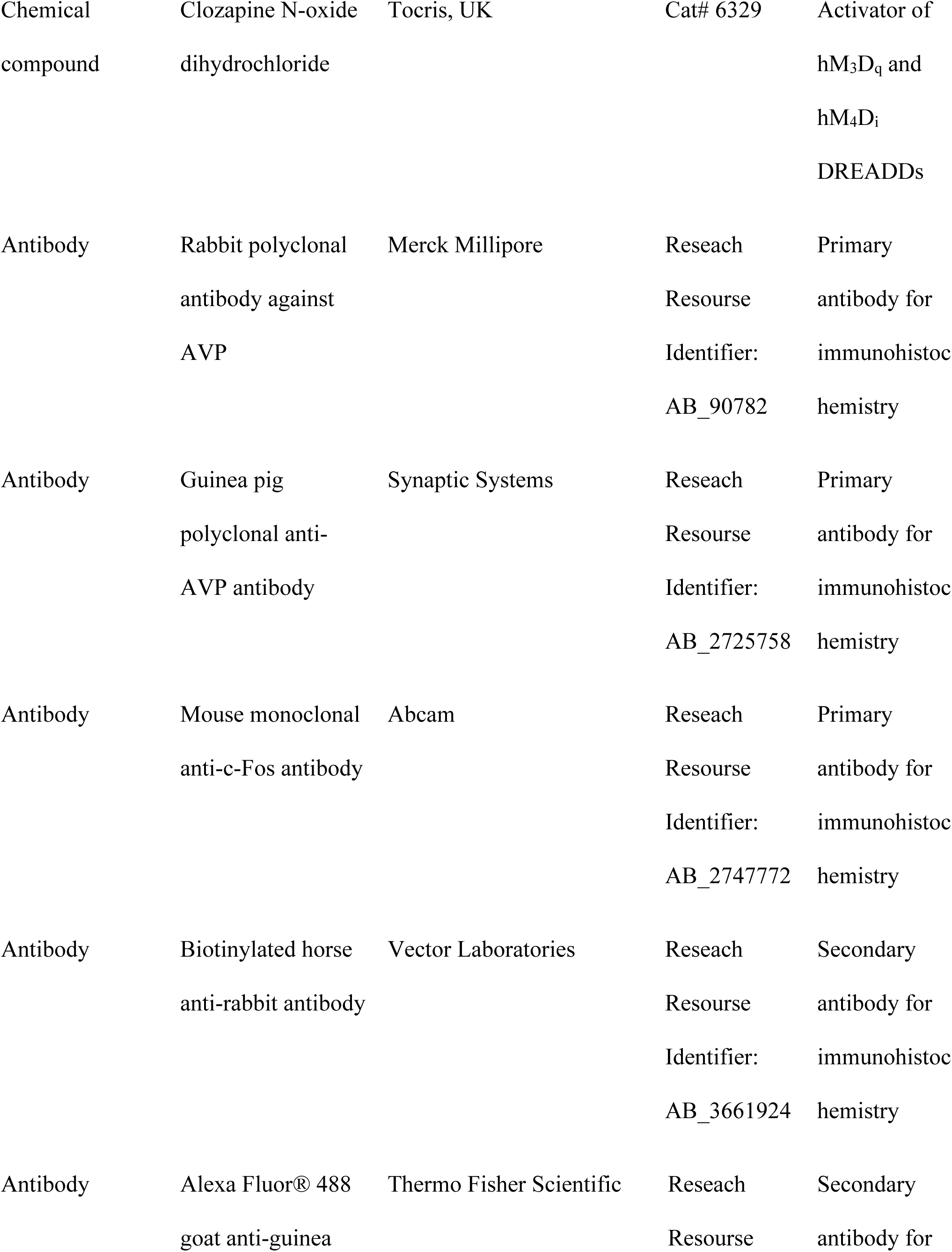
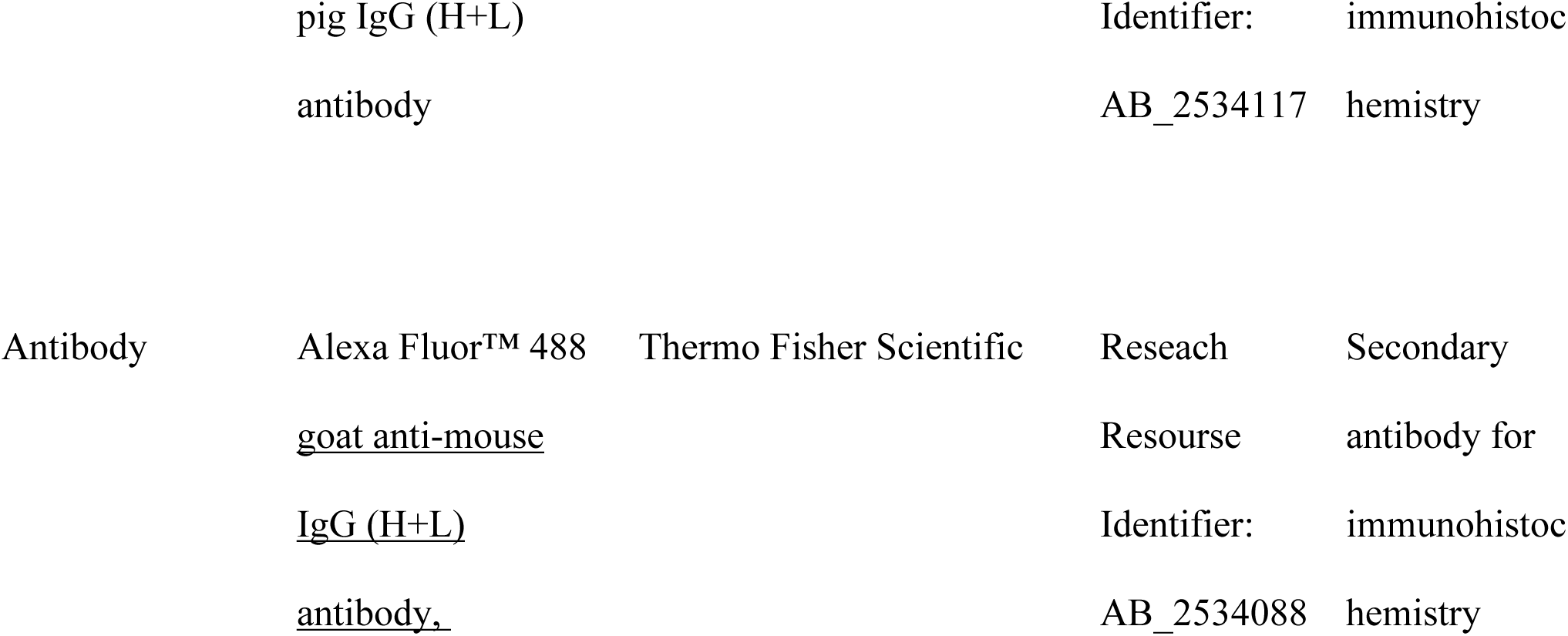

